# Hydroxyurea induces ER stress and cytoplasmic protein aggregation

**DOI:** 10.1101/2024.10.17.618805

**Authors:** Ana Sánchez-Molina, Manuel Bernal, Joel D. Posligua-García, Antonio J. Pérez-Pulido, Laura de Cubas, Elena Hidalgo, Silvia Salas-Pino, Rafael R. Daga

**Affiliations:** Centro Andaluz de Biología del Desarrollo, Univ. Pablo de Olavide, Seville, Spain; Departamento de Biología Molecular y Bioquímica, Facultad de Ciencias, Universidad de Málaga, Andalucía Tech, 29071 Málaga, Spain; Instituto de Investigación Biomédica de Málaga y Plataforma en Nanomedicina, IBIMA Plataforma BIONAND, 29590 Málaga, Spain; Oxidative Stress and Cell Cycle Group, Universitat Pompeu Fabra, Barcelona, Spain

**Keywords:** Hydroxyurea, Thiol Stress, Diamide, Glutathione, Endoplasmic Reticulum, Endoplasmic Reticulum Expansion, Nuclear Pore Complex, Nuclear Architecture, Oxidative Stress, Protein Folding, Protein Aggregation

## Abstract

The endoplasmic reticulum (ER) lumen provides the proper redox environment for disulfide bond formation, which is required for the appropriate folding of proteins that enter the secretory pathway and constitute membranes. Defective protein folding in the ER activates proteostatic mechanisms that are now beginning to be elucidated. Here, we show that hydroxyurea (HU) causes ER stress and triggers a transient perinuclear ER expansion, which leads to the clustering of nuclear pore complexes. This striking phenotype is mimicked by diamide (DIA), a specific thiol stress inductor, and prevented or rapidly reverted by dithiothreitol, a dithiol-reducing agent, suggesting that ER expansion is caused by disulfide stress. ER expansion induced by HU or DIA depends on glutathione (GSH), is Ire1-independent, and is associated with a unique transcriptome program that differs from the canonical unfolding protein response (UPR). The ER luminal expansion accumulates Hsp70 Bip1 chaperone, and it evolves parallel with the appearance of cytoplasmic protein aggregates containing heat stress proteins (HSPs), indicating that both HU and DIA are impinging on protein folding. Thus, our data reveal that HU induces disulfide stress that impinges on protein folding in the cytoplasm and ER.

## INTRODUCTION

The endoplasmic reticulum (ER) is the largest endomembrane organelle in the cell. It consists of a complex network of tubules and membranes that are continuous with the outer nuclear membrane (ONM), from where the ER expands into the cytoplasm and physically connects with the plasma membrane^1,2^. The inner nuclear (INM) and the ONM/ER form a unique, continuous membrane (perinuclear ER) that bends at the nuclear pore complexes (NPCs)^3,4^.

The ER is responsible for the synthesis, folding, and assembly of secretory and membrane proteins, which represent almost one-third of the eukaryotic proteome^5,6^. To achieve their functional conformation, nascent proteins undergo a sequential folding process in the ER lumen, coordinated with post-translational modifications such as glycosylation and disulfide bond formation^5,7–9^. Defects in redox folding can lead to protein misfolding and aggregation, causing both ER and cellular stress^10^. To prevent this, cells monitor these processes through the coordinated action of protein disulfide isomerases (PDIs) and ER oxidoreductases^11,12^. When proteins fail to fold correctly in the ER, cells activate the unfolded protein response (UPR), a coordinated network of signaling cascades which collaboratively, reduce overall mRNA translation, increase the folding capacity of the ER and contribute to the removal of unfolded proteins through ER-Associated Degradation (ERAD) and autophagy^13,14^. Additionally, in response to ER stress, the UPR activates lipid biosynthesis, prompting membrane expansion and subsequently increasing ER size, which helps mitigate folding stress^15^. In cases where stress is unresolved, UPR eventually prompts cell death^16^.

Glutathione (GSH) plays a central role in redox regulation, as it counters the oxidation potential by maintaining a significant fraction of PDIs and oxidoreductases in the reduced state, and directly contributes to the reduction of disulfides along with the thioredoxin and glutaredoxin systems^17–23^. GSH also plays an important role in the biosynthesis and maturation of Iron-Sulfur (Fe-S) clusters in proteins^22,24^. In the ER, the GSH/oxidized glutathione (GSSG) ratio is around 1.5:1 to 3.3:1^25^ to facilitate disulfide bond formation and proper oxidative protein folding. In contrast, in the cytosol, disulfide bonds rarely form due to the high GSH/GSSG ratio (∼100:1)^19,26,27^, which favors proteins to remain in their reduced state^28^. Imbalances in the cytosolic GSH/GSSG ratio can alter global GSH/GSSG levels^29^, which in turn may lead to impairments in mitochondrial redox regulation^24,30^.

In this work, we show that two prooxidant molecules, hydroxyurea (HU), an inhibitor of ribonucleotide reductase, and diamide (DIA), a specific thiol stress inductor, cause thiol stress in the ER. The thiol-specific ER stress response triggers a transient phenotype of perinuclear ER expansion involving nuclear envelop dilation that leads to the clustering of the whole population of NPCs, along with the spindle pole body (SPB), the yeast equivalent of the centrosome, in one region of the NE. ER expansion induced by thiol stress accumulates ER specific Hsp70 chaperone Bip1 while several other HSPs accumulate in cytoplasmic aggregation foci. Our results suggest that HU induces disulfide stress that compromises proper protein folding in the ER and cytoplasm and elicits proteostatic mechanisms in both compartments.

## RESULTS

### HU causes NPC clustering and ER expansion independent of RNR activity inhibition

HU is a potent and reversible inhibitor of ribonucleotide reductase (RNR) used as an FDA-approved antiproliferative drug in cancer therapy. In addition, HU is the primary treatment for sickle cell anemia, is used as an antiviral, and to treat several infectious diseases^31^. In research laboratories, HU is commonly used to synchronize cells in S phase. By doing so, we unexpectedly found that HU treatment in *S. pombe* resulted in an altered localization of NPCs. Incubation of cells expressing nucleoporin Cut11-GFP (a component of the NPC membrane ring) with 15 mM HU for four hours, a concentration and time known to arrest cell cycle in S phase^32,33^, disrupted the even localization of NPCs and resulted in their accumulation along with the SPB (as shown with Sid2-Tom) in a localized region of the NE (Figure 1A-B). This previously undescribed phenotype was dose-dependent, as increasing the concentration of HU resulted in up to 97.33±2.08% (n>100 cells) of cells showing NPC clustering (Figure 1A, graph).

**Figure 1.**
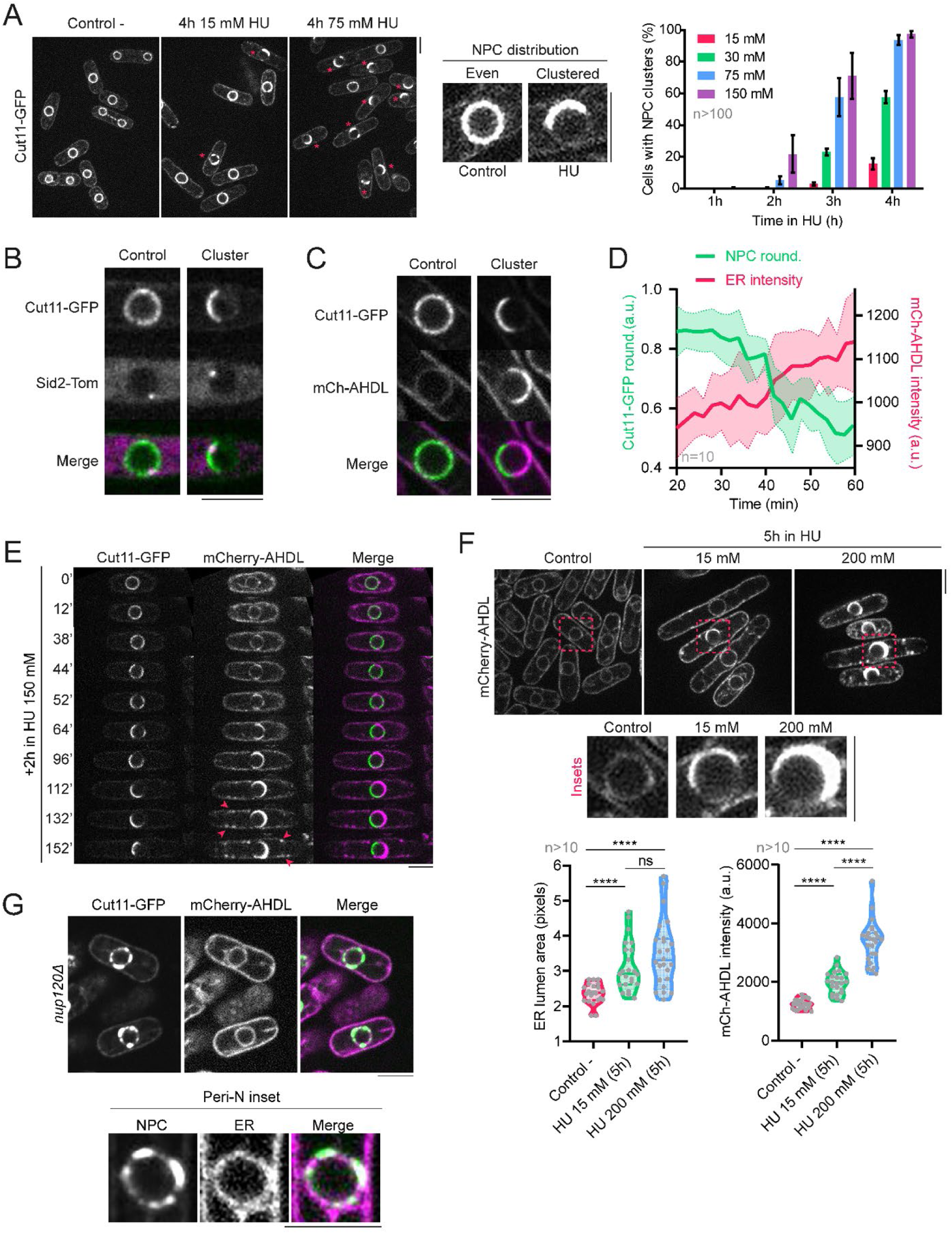
Hydroxyurea induces NPC clustering and ER expansion. **(A) Left panels:** Confocal microscopy images of wild-type cells with GFP-tagged nucleoporin Cut11 under control untreated conditions (left) and after 4 hours at 30°C in 15 mM HU (center) or 75 mM HU (right). Magenta asterisks marks cells in which NPC clustering can be observed. **Central panels:** Insets showing a close-up of the even distribution of NPCs observed in an untreated cell versus the phenotype of clustered NPCs observed in HU. **Right panel:** Quantification of the percentage of cells showing clustered NPCs after treatment with different concentrations of HU for several hours. The graph shows the mean ± SD of three independent repetitions of the experiment, and in each repetition n>100 cells were accounted for each HU concentration, temperature and time point. **(B)** Nuclear insets of confocal microscopy images showing a control nucleus expressing the SPB marker Sid2-tomato and in which Cut11-GFP is evenly distributed along the NE, and a nucleus where NPCs cluster in one region of the NE and the SPB localizes in that same region after HU treatment. **(C)** Nuclear insets of confocal microscopy images showing a control nucleus in which Cut11-GFP and the luminal ER marker mCherry-AHDL colocalize along the NE, and a nucleus where Cut11-GFP and mCherry-AHDL occupy opposite sides of the nuclear periphery after HU treatment. **(D)** Quantification of the loss of roundness of NPCs along the NE (from 1, full roundness, to 0.4, less than half of the circumference of the NE decorated with NPCs), which correlates with an increment in mCherry-AHDL fluorescence intensity (measured in arbitrary units) along the nuclear periphery. Graph shows the mean ± SD of ten representative cells which develop the phenotype along the course of the experiment. **(E)** Timelapse showing confocal microscopy images depicting NPC clustering and ER expansion in a cell expressing Cut11-GFP and mCherry-AHDL after exposure to 150 mM HU for two hours before the beginning of the experiment. Arrows show the appearance of cortical ER expansions. **(F) Upper panels:** confocal microscopy images showing mCherry-AHDL to compare luminal ER size and intensity in a control untreated condition and after exposure to 15 mM and 200 mM HU for 5 hours. Insets show a close-up of a significant nucleus in each condition. **Lower panels:** Quantifications of ER area, measured in pixels, and mCherry-AHDL fluorescent intensity in the nuclear periphery after a 5-hour incubation in either 15 mM or 200 mM HU, compared to the untreated condition. Graphs show violin plots with their respective mean ± SD, where at least 10 cells were accounted for in each condition. **(G)** Confocal microscopy images and nuclear insets of the mutant strain *nup120Δ*, where NPCs tagged with Cut11-GFP are not evenly distributed along the NE while the luminal ER marker mCherry-AHDL remains unaltered in the nuclear periphery. All confocal microscopy images are SUM projections of three central Z slices. Scale bars represent 5 microns.

During HU treatment, clustered NPCs remain functional for nucleocytoplasmic transport. This was demonstrated by measuring the rate of importin-α (Imp1) transport across evenly distributed NPCs and clustered NPCs in HU-exposed cells by fluorescence recovery after photobleaching (FRAP), bleaching either the cytoplasmic or nuclear pools of Imp1-GFP, and analyzing the fluorescence recovery in the adjacent compartment (Figure S1A). We also determined that the translocation rate across NPCs of the transcription factor Pap1 upon H_2_O_2_ addition, which induces its translocation to the nucleus, and after H_2_O_2_ washout, which induces Pap1 transport back to the cytoplasm^34–37^, was similar in cells with normal NPCs and cells with clustered NPCs in the presence of HU (Figure S1B).

**Supplementary figure 1.**
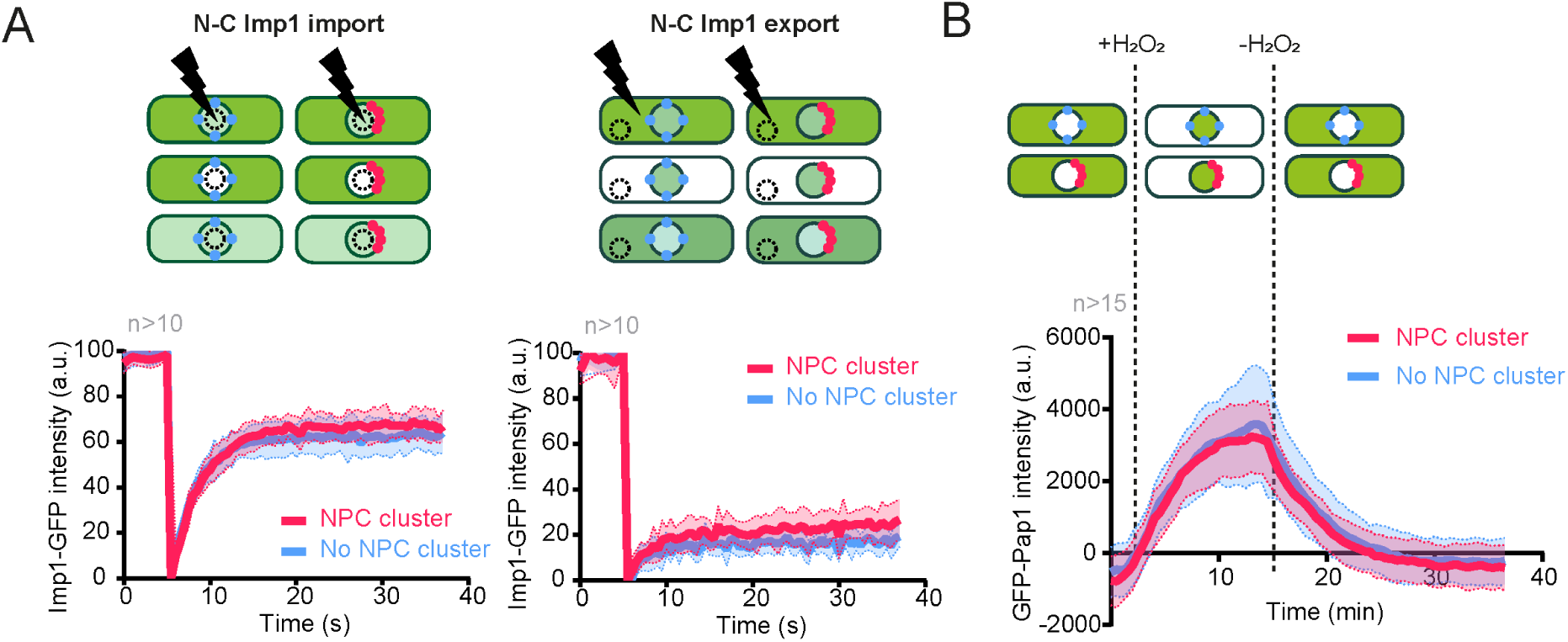
NPCs clustered after HU treatment remain functional for nucleocytoplasmic transport. **(A)** Recovery of importin (Imp1-GFP) fluorescence after FRAP in the nucleus (left) and in the cytoplasm (right) in cells with and without clustered NPCs, both populations exposed to 15 mM HU for 4 hours. Graphs represent the mean ± SD of the fluorescence intensity of Imp1-GFP, normalized to the fluorescence in the indicated compartment right before bleaching, and measured in at least 10 cells of each phenotype. **(B)** Progression of GFP-Pap1 intensity after exposure and removal of 0.2 mM H2O2, in cells with and without clustered NPCs, after a 4-hour exposure to 15 mM HU. Graph represents the mean ± SD of the fluorescence intensity of GFP-Pap1, normalized to the background and measured in at least 15 cells of each phenotype.

As the primary target of HU is RNR, we checked whether NPC clustering was related to its inhibition or to the cell cycle arrest in S phase. RNR is a highly conserved tetrameric enzyme composed of two large subunits (Cdc22^R1^) and two small subunits (Suc22^R2^). Whereas Cdc22^R1^, which bears the catalytic activity, is mainly cytoplasmic, Suc22^R2^ is actively imported into the nucleus in a manner dependent on the regulator Spd1. Upon HU treatment, Spd1 is degraded, allowing cytoplasmic accumulation of Suc22^R2^ ^38,39^. We determined that the appearance of clustered NPCs was unrelated to the differential nucleocytoplasmic compartmentalization of the two RNR subunits, as deletion of *spd1*, which results in both subunits constitutively localizing in the cytoplasm^39^, did not alter the frequency of NPC clustering (Figure S2A). Inactivation of Cdc22^R1^ by using a thermosensitive *cdc22-M45* mutant strain and keeping it at restrictive temperature for over four hours^40,41^ did not result in NPC clustering (Figure S2B). Furthermore, HU-dependent NPC clustering could be induced in G2/M blocked cells (by inactivation of *cdc25-22*, a thermosensitive allele of the mitotic activator Cdc25 (Figure S2C)). Together, these results indicate that HU-induced NPC clustering is a phenomenon not related to S phase arrest nor to deoxyribonucleotide synthesis inhibition, which is the main known cellular effect of HU, and instead is the result of an undescribed off-target of this drug.

**Supplementary figure 2.**
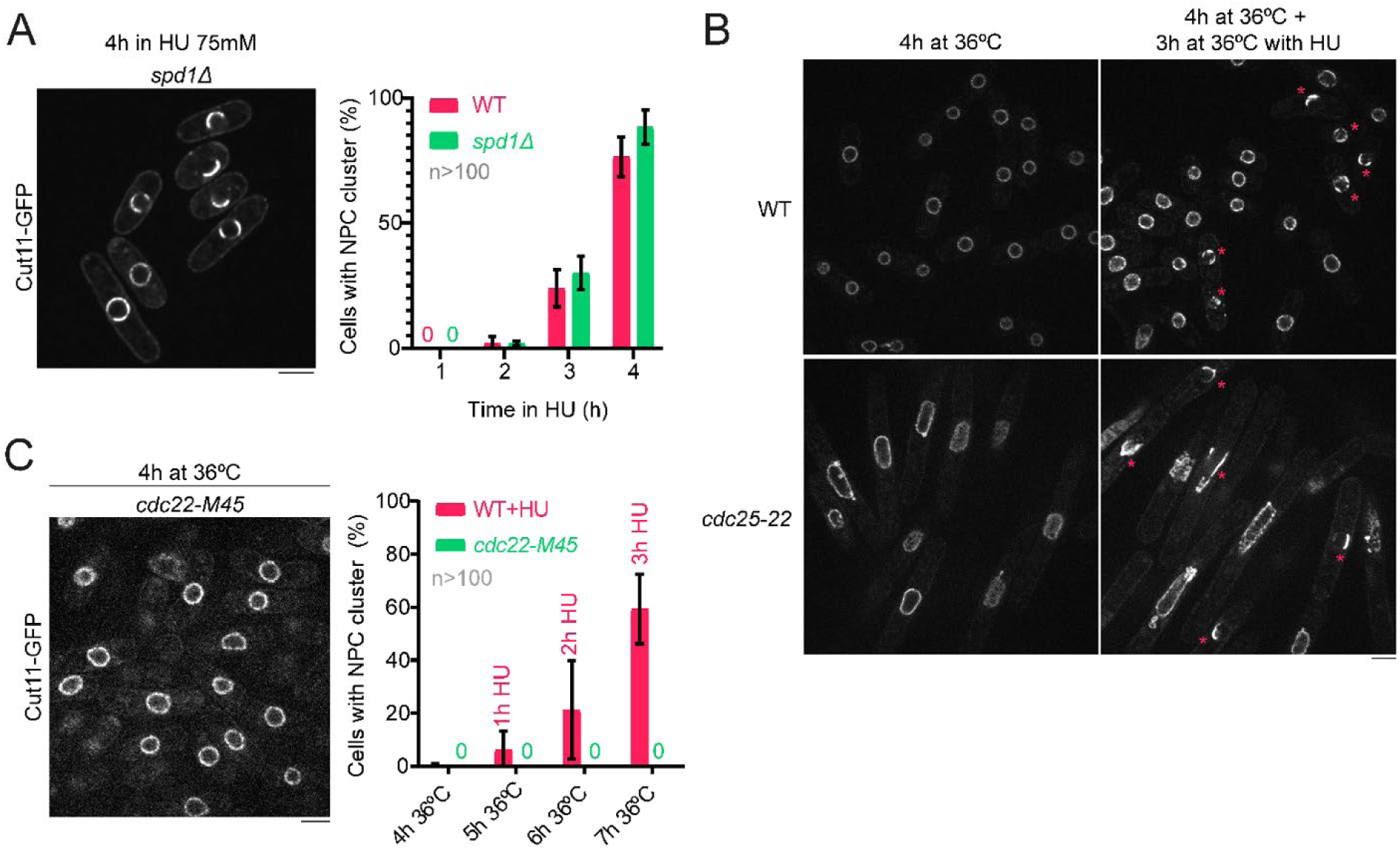
NPC cluster formation is independent of S phase arrest induced by RNR inhibition. **(A)** Representative confocal microscopy image of a *spd1Δ* mutant strain after 4 hours in 75 mM HU showing clustered NPCs (left), and graph comparing NPC cluster formation in a wild-type strain and a *spd1Δ* mutant (right). Images are SUM projections of three central Z slices. Scale bars represent 5 microns. Graph shows the mean ± SD of two independent repetitions of the experiment, and in each repetition at least 100 cells were accounted for each condition. **(B) Left:** Representative confocal microscopy image of a *cdc22-M45* thermosensitive mutant after 4 hours at restrictive temperature (36°C) proving that these cells do not form NPC clusters *per se* (left), and graph comparing NPC cluster formation in a control wild-type strain exposed to 75 mM HU at 37°C and a *cdc22-M45* mutant kept at 36°C for 4 hours and then exposed to 75 mM HU for the following 3 hours while still at restrictive temperature (right). Images are SUM projections of three central Z slices. Scale bars represent 5 microns. Graph shows the mean ± SD of two independent repetitions of the experiment, and in each repetition at least 100 cells were accounted for each condition. **(C)** Representative confocal microscopy images of a wild-type and a *cdc25-22* thermosensitive mutant, both expressing Cut11-GFP, at restrictive temperature (36°C) after incubation for 4 hours (left) to achieve full blockage of the mutant in the G2/M transition, and after addition of HU for 3 extra hours while kept at 36°C so cell cycle in the mutant remains blocked (right). When *cdc25-22* cells are blocked in G2/M transition by temperature shift and treated with HU during this blockage, cells form NPC clusters while in G2/M. Magenta asterisks indicate cells with NPC clustering. Images are SUM projections of five central Z slices. Scale bars represent 5 microns.

We noticed that NPC clustering was accompanied by a severe asymmetry in the perinuclear ER morphology, in which the perinuclear ER luminal space (marked with an mCherry tag fused to the AHDL localization signal^42,43^) entirely occupied the space devoid of NPCs (Figure 1C). Timelapse confocal microscopy images showed that NPC clustering and perinuclear ER lumen enlargement occurred concomitantly (Figure 1D-E). After the initial expansion of the perinuclear ER lumen, events of cortical ER expansion were also observed when HU exposure time increased (Figure 1E, arrows). Similar to NPC clustering, HU-induced perinuclear ER enlargement was time- and dose-dependent (Figure 1F). Thus, HU induces an overall change in nuclear architecture, including expansion of perinuclear luminal ER and NPC clustering, a phenotype that we named ‘Nuclear Cap’ (N-Cap).

We next asked whether enlarged perinuclear ER resulted from the clustering of NPCs or if, on the contrary, remodeling of the perinuclear ER was causing NPCs to cluster. For that, we used a strain deleted for the nucleoporin Nup120, a component of Y-complex of the inner and outer NPC rings and whose absence results in defective and aberrantly clustered NPCs^44^. Clustered NPCs in *nup120Δ* cells showed an apparently normal perinuclear ER morphology (Figure 1G), indicating that NPC clustering *per se* does not cause detectable perinuclear ER asymmetry. Therefore, we concluded that the HU-induced N-Cap phenotype consists of an enlargement of the perinuclear ER lumen, which likely triggers the clustering of NPCs in one region of the NE opposite the expanded ER.

### Perinuclear ER expansion involves INM and ONM/ER separation

To further characterize the phenotype of perinuclear ER expansion, we analyzed by live cell microscopy the localization of INM proteins Lem2, Man1 and Ima1, as well as of Cut11, which marks both NPCs and NE, in cells with expanded ER after exposure to DIA for 4 hours (Figure S3A). Cut11 localized at clustered NPCs (Figure 1B-C), although it was also detected at the INM and the expanded ER. Man1 also accumulated coincident with the localization of NPCs, which is consistent with recent findings^45^; however, Lem2 and Ima1 were enriched at the INM, although a faint signal appeared also at the expanded ER.

We further performed transmission electron microscopy (TEM) of HU-treated cells in the conditions of maximum ER expansion (incubation for 4 hours) along with wild-type untreated cells. TEM images revealed perinuclear ER expansion and lumen enlargement with INM and ONM/ER membrane separation (Figure S3B-C). Within the enlarged luminal ER, globular structures could be detected, which might represent protein aggregates or misfolded peptides (Figure S3C, black arrow). Thus, both live cell images of GFP-tagged INM proteins and TEM images revealed that the perinuclear ER expansion involves separation of the INM and ONM, likely due to the accumulation of misfolded peptides, which, in turn, constricts NPCs in the opposite region of the NE.

**Supplementary figure 3.**
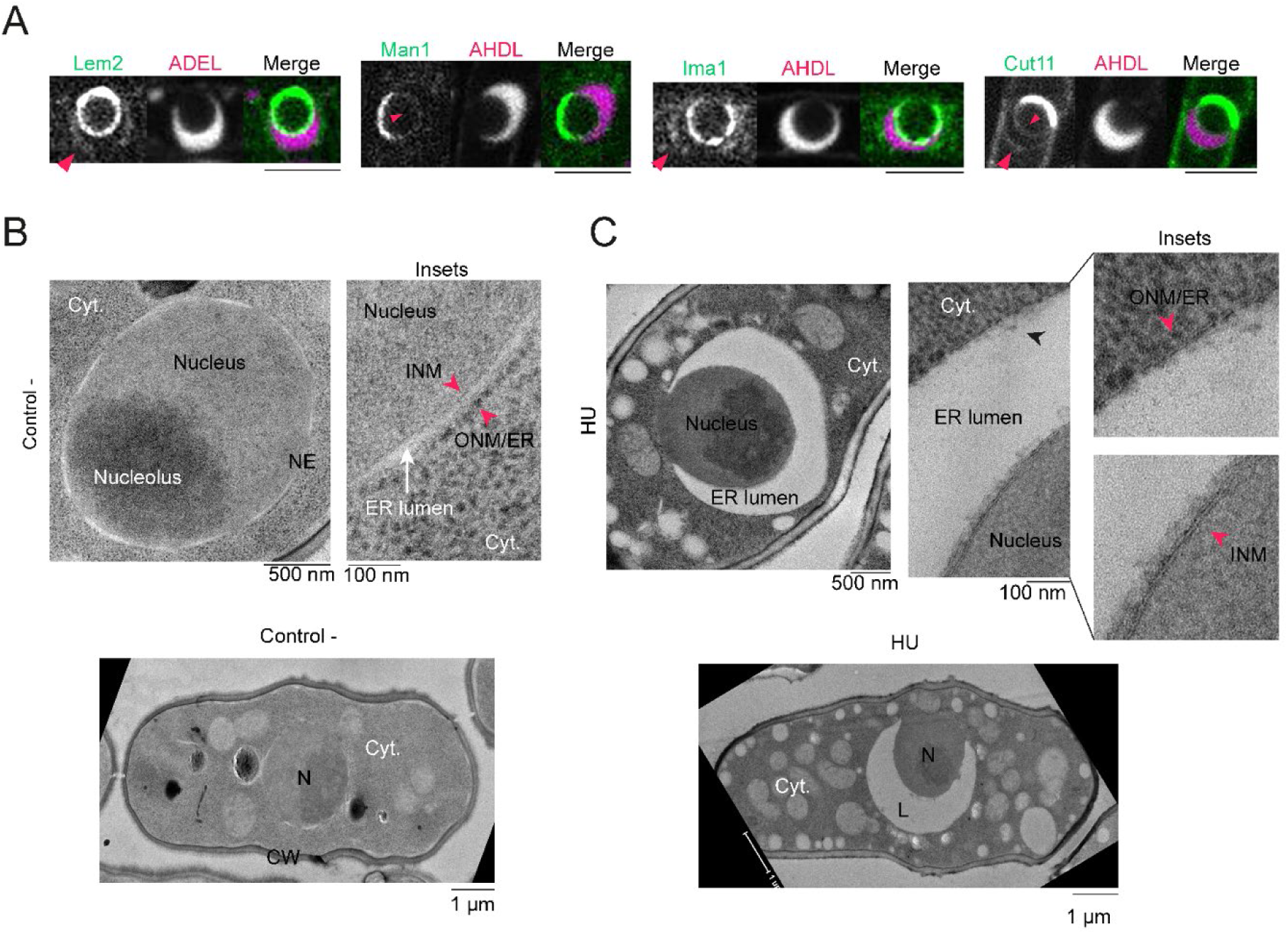
HU- and DIA-induced ER expansion causes INM and ONM/ER membrane separation. **(A)** Confocal microscopy images of cells expressing INM tags Lem2-GFP, Man1-GFP and GFP-Ima1, or NPC tag Cut11-GFP, opposed to the luminal ER marker mCherry-AHDL, after a 4-hour incubation in 75 mM HU. Magenta arrows show the presence of membranes surrounding the ER lumen. Images are SUM projections of three central Z slices. Scale bars represent 5 microns. **(B-C)** Transmission electron images of a wild-type strain in control conditions (B) and after a 4-hour incubation in 75 mM HU (C). Insets focus on nuclei in each condition; magenta arrows indicate inner nuclear membranes (INM) and outer nuclear membranes continuous with the ER (ONM/ER); black arrows indicate electrodense formations found in the expanded ER after DIA exposure. N = nucleus; CW = cell wall; NE = nuclear envelope; Cyt. = cytosol. Scale bars represent 1 micron, 500 nm or 100 nm, as indicated in each panel.

### HU-induced perinuclear ER expansion and NPC clustering are reversible phenotypes

To determine whether HU-induced N-Caps were a deleterious and terminal phenotype or a transient response to cellular damage, we induced their formation in cells expressing Cut11-GFP and mCherry-AHDL by incubation in 75 mM HU-containing media for 3 hours. Then, HU was washed out and cells were filmed by confocal microscopy. We found that both the perinuclear luminal ER and NPCs concomitantly recovered their isometrical distribution along the nuclear periphery on average 80.86±31.31 (n=37) minutes following HU washout (Figure 2A-B, S4A-B). Interestingly, N-Cap dissolution showed a biphasic kinetics, with a first long ‘lagging’ phase followed by a rapid recovery phase in which perinuclear architecture returned to a distribution typical of unperturbed cells (Figure S4A-B). ER restoration shortly preceded NPC redistribution, supporting the hypothesis that ER expansion triggers NPC clustering (Figure S4B, black arrow). Importantly, if HU treatment is maintained for a longer time, perinuclear ER and clustered NPCs eventually returned to their normal architecture (Figure 2C) indicating that cells adapt to high doses of HU. We also found that the time needed to restore normal NPC distribution depended on the size of the perinuclear ER lumen, as cells with a larger ER lumen took longer to redistribute their NPCs and restore their ER in comparison to cells that had a less expanded ER in the same conditions (Figure S4C-D). Therefore, our results suggest that HU induces ER stress manifested as a transient enlargement of perinuclear ER lumen.

**Figure 2.**
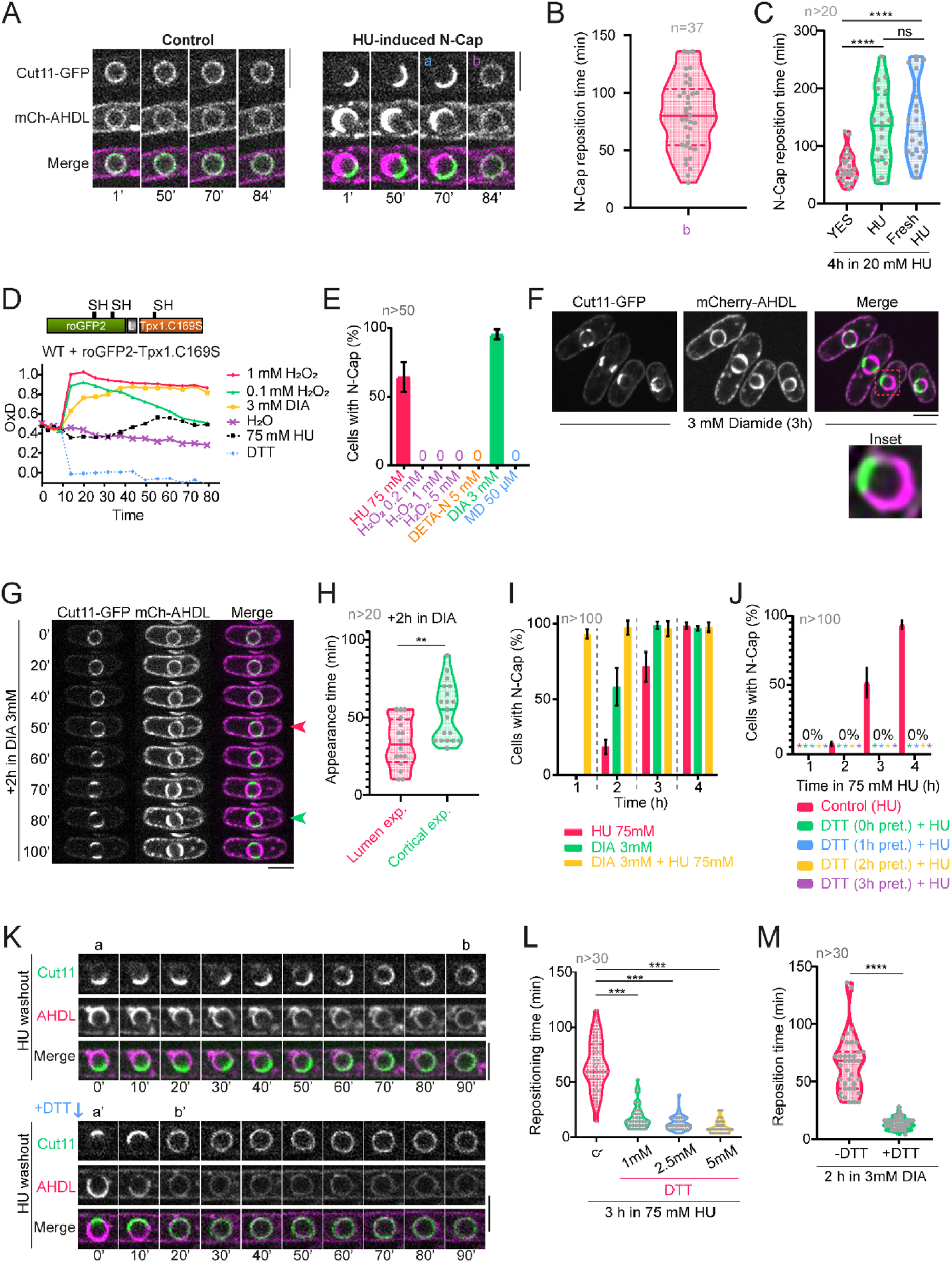
HU-induced ER expansion is reversible and mimicked by thiol stress. **(A)** Confocal microscopy images comparing a nucleus with even NPC and ER distributions, as tagged by Cut11-GFP and mCherry-AHDL respectively (left panels), and a nucleus with an N-Cap after incubating 3 hours in 75 mM HU (left panels). ‘a’ marks the last timepoint before N-Cap redistribution; ‘b’ marks the recovery of the even perinuclear architecture. **(B)** Quantification of the time N-Caps take to recover their isometric distribution (‘b’) along the NE after a 3-hour incubation in 75 mM HU following drug washout. Graph shows a violin plot with the mean ± SD of the 37 cells accounted for in the experiment. **(C)** Quantification N-Cap reposition time after a 4-hour incubation in 20 mM HU. ‘YES’ indicates that HU was washed out of the medium and clean medium was inoculated to the cells; ‘HU’ indicates that the medium remains the same from the start of the incubation with HU; ‘Fresh HU’ indicates that fresh YES media with HU was inoculated to the cells at the same time cells under the ‘YES’ category were inoculated with clean medium. Graph shows violin plots with the mean ± SD of the reposition time of at least 20 cells accounted for in each condition. **(D)** H_2_O_2_ production by HU using the biosensor roGFP2-Tpx1.C169S. Quantification of the degree of probe oxidation, or OxD (amount of probe oxidation per 1), of roGFP2-Tpx1.C169S over time. Cultures of strain HM123 carrying plasmid p407.C169S were treated with 1 mM H_2_O_2_ (to achieve maximal probe oxidation, 1), 50 mM DTT (it induces full reduction of the probe, 0), water (control), 0.1 mM H_2_O_2_, 75 mM HU or 3 mM DIA. The average of three biological replicates is represented. **(E)** Quantification of the incidence of the N-Cap phenotype in 75 mM HU, 0.2 mM H_2_O_2_, 1 mM H_2_O_2_, 5 mM H_2_O_2_, 5 mM DETA-NONOate (DETA-N), 3 mM DIA and 50 μm menadione (MD) after a 3-hour incubation. Apart from HU, only DIA reproduces the N-Cap phenotype. **(F)** Confocal microscopy images of cells treated with 3 mM DIA for 3 hours and an inset of a representative nucleus showing the N-Cap phenotype, as seen with tags Cut11-GFP and mCherry-AHDL. **(G)** Confocal microscopy images showing a timelapse of the cellular architecture modification after DIA exposure. The magenta arrow marks N-Cap apparition; the green arrow marks the apparition of cortical ER expansions. **(H)** Quantification of the apparition time of the N-Cap phenotype (‘Lumen exp.’ stands for ‘lumen expansion’) and of the cortical expansions (‘Cortical exp.’). Graph shows violin plots with the mean ± SD of the apparition time of the indicated phenotype of at least 20 cells. **(I)** Quantification of the incidence of the N-Cap phenotype in a population of cells over time in 75 mM HU, 3 mM DIA or a combination of the two. Combining HU and DIA leads to a synergic effect upon N-Cap formation. **(J)** Quantification of the incidence of the N-Cap phenotype in a cell population during a 4-hour treatment with 75 mM HU (‘Control (HU)’), during a treatment combining 75 mM HU and 1 mM DTT (‘DTT (0h pret.)+HU’), or during the same treatment with a prior pretreatment with 1 mM DTT for 1 hour (‘DTT (1h pret.)+HU’), 2 hours (‘DTT (2h pret.)+HU’) or 3 hours (‘DTT (3h pret.)+HU’). **(K)** Confocal microscopy images comparing a timelapse of N-Cap reposition in two representative nuclei after a 3-hour incubation in 75 mM HU, one in which clean YES medium is added (upper panel) and the other with 1 mM DTT added when HU is washed out of the medium (lower panel). Addition of DTT to the medium when washing HU from a culture leads to the rapid redistribution of NPCs and ER, as seen with tags Cut11-GFP and mCherry-AHDL respectively. **(L)** Quantification of the reposition time of N-Caps after HU washout. Addition of DTT during HU washout significantly reduces the time it takes for cells to reposition their internal architecture after a 3-hour incubation in 75 mM HU. Higher DTT concentrations cause a faster dispersion of the phenotype. **(M)** Quantification of the reposition time of N-Caps after DIA washout with and without DTT added to the medium after a 2-hour incubation in 3 mM DIA. N-Caps formed in DIA dissolve when the drug is washed out from the medium, and this dissolution occurs faster when adding DTT. All confocal microscopy images are SUM projections of three central Z slices. Scale bars represent 5 microns. Graphs in (B), (L) and (M) show violin plots with the mean ± SD of the reposition time of N-Caps in least 20-30 cells (as indicated in the n) accounted in each condition. Graphs in (E), (I) and (J) show the mean ± SD of two independent repetitions of each experiment, with at least 50-100 cells (as indicated in the n) accounted in each condition. See Figure S4 for more information.

**Supplementary figure 4.**
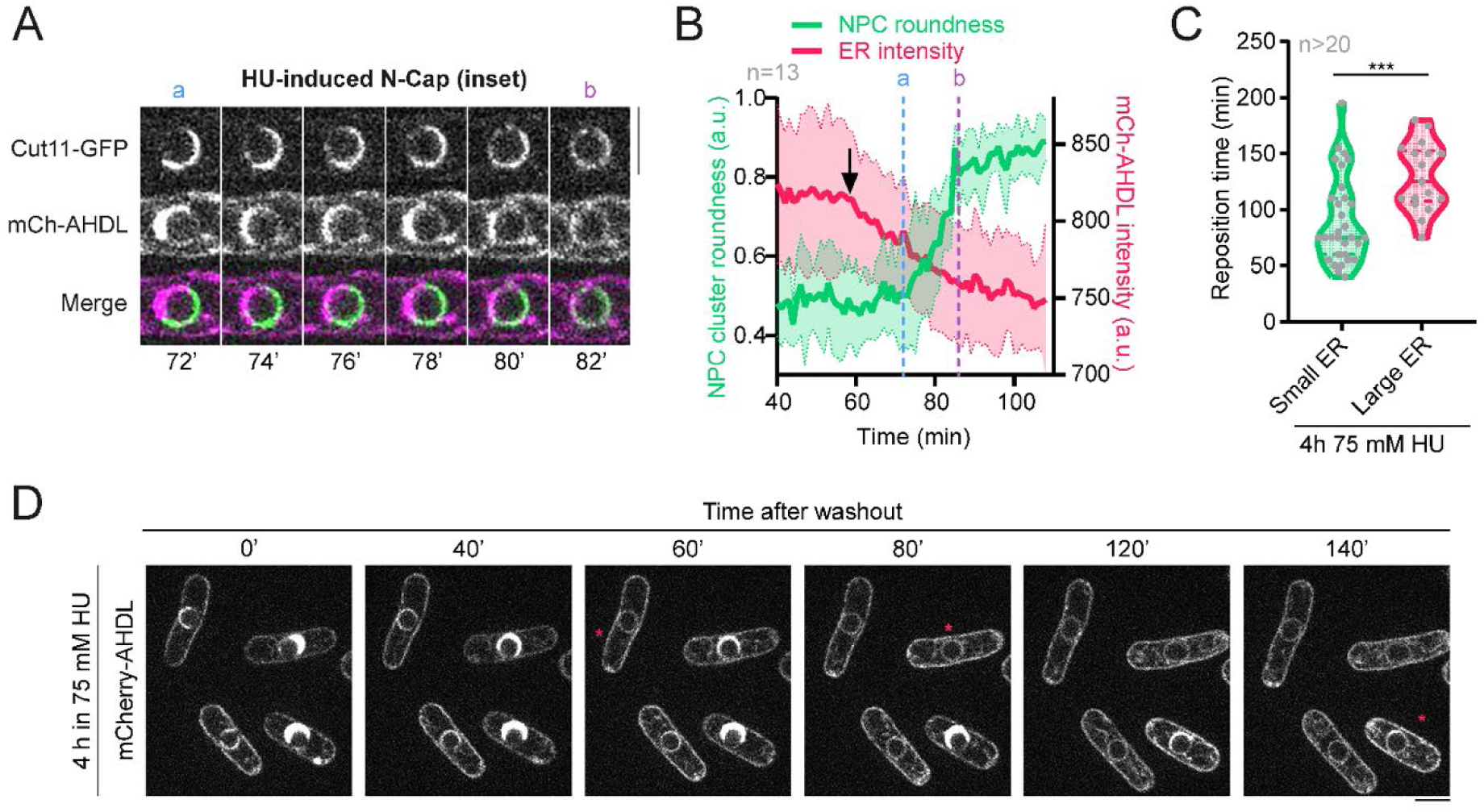
Dispersion of NPC clusters require perinuclear ER redistribution. **(A)** Insets of confocal microscopy images showing the reposition of the N-Cap indicated between ‘a’ and ‘b’ in Figure 2A. Images are SUM projections of three central Z slices. Scale bars represent 5 microns. **(B)** Progression of the roundness of the NPCs during N-Cap reposition compared to mCherry-AHDL intensity. The black arrow shows the point where the perinuclear ER intensity starts to decay, which occurs before the rapid redistribution of NPCs that takes place between ‘a’ and ‘b’. Graph shows the mean ± SD of 13 representative cells that reposition their perinuclear architecture along the course of the experiment. **(C)** Quantification of N-Cap reposition time after HU washout following a 4-hour incubation in 75 mM HU, comparing cells with a more expanded ER (‘large ER’) versus cells with a less expanded ER (‘small ER’). Graph shows violin plots with the mean ± SD of the reposition time of N-Caps of at least 20 cells with each of the forementioned perinuclear ER morphologies. **(D)** Timelapse of confocal microscopy images following a group of cells tagged with mCherry-AHDL, previously exposed to 75 mM HU for 4 hours, after drug washout. Cells with a larger ER take longer to recover the even perinuclear architecture than those with a less-expanded ER. Asterisks mark when a cell recovers its normal perinuclear architecture. Images are SUM projections of three central Z slices. Scale bars represent 5 microns.

### HU-induced perinuclear ER expansion is mimicked by thiol stress

It has been shown that HU treatment produces reactive oxygen species (ROS) *in vivo*, including hydrogen peroxide (H_2_O_2_) and nitric oxide (NO)^46,47^. We confirmed that HU induces oxidative stress in fission yeast, as the biosensor roGFP2-Tpx1.C169S, which is finetuned for detecting H_2_O_2_ fluctuations, showed a clear oxidation in the cytosol during HU treatment, similar to oxidants diamide (DIA) or H_2_O_2_, although H_2_O_2_ and specially DIA oxidize the sensor faster and more effectively (Figure 2D). Thus, we checked whether the HU-induced ER expansion was due to oxidative stress produced by H_2_O_2_, which is a byproduct of HU oxidation. We found that treatment with increasing concentrations of H_2_O_2_ did not induce N-Cap formation (Figure 2E). Addition of DETA-NONOate, a compound that generates NO, did not produce any observable ER phenotype either (Figure 2E). We further tested if treatment with other oxidants, such as menadione (MD), which generates superoxide^48^, and DIA, which is a membrane-permeable specific thiol oxidant^49,50^, resulted in N-Cap formation. Whereas MD did not produce any noticeable N-Cap phenotype (Figure 2E), DIA mimicked the effect of HU and led to the emergence of N-Caps that were indistinguishable from those observed during HU treatment (Figure 2E-F). Similarly to HU, both NPC clustering and perinuclear ER lumen expansion emerged concomitantly during DIA treatment, and DIA also induced cortical ER expansion at longer exposition times, once the perinuclear ER had already expanded and the N-Cap phenotype had emerged (Figure 2G-H). Furthermore, DIA and HU acted synergistically, vastly increasing the incidence of N-Caps when combined (Figure 2I).

DIA has been shown to oxidize GSH to GSSG, thus depleting reduced glutathione in the cell^49^ and indirectly promoting disulfide bond formation and glutathionylation in proteins containing thiol groups (-SH), which is known as thiol or disulfide stress^51,52^. Conversely, dithiothreitol (DTT) is a strong reductant that turns oxidized cysteines back to their reduced state, breaks disulfide bonds and reverts protein glutathionylation^53,54^. Thus, we hypothesized that DTT treatment might prevent and/or revert N-Cap formation if this HU-induced phenotype was due to hyperoxidation of thiol groups, as suggested by the reproduction of this phenotype in DIA. Consistently, when cells were pretreated with DTT before HU addition, the appearance of N-Caps was fully suppressed (Figure 2J). Moreover, DTT addition to HU- or DIA-treated cells displaying N-Caps reverted this phenotype with accelerated kinetics than simple drug washout (Figure 2K-M). Thus, these data indicate that perinuclear ER expansion induced by HU or DIA is likely caused by disulfide stress.

Finally, we addressed whether HU-induced NPC clustering and ER expansion were evolutionarily conserved. For that, we first tested if NPC clustering was observed in the budding yeast *S. cerevisiae* and, indeed, in cells expressing NUP84-mCherry, NPC clusters were noticed after a 3-hour incubation in 100 mM HU (Figure S5A) and the dynamic NPC reposition after HU washout was also recapitulated (Figure S5B), indicating that this phenotype is not specific to *S. pombe*. We further confirmed the conservation of the ER expansion phenotype in mammalian cells by using HT1080 fibrosarcoma cells and incubating them with 200 µM and 1 mM HU (Figure S5C). ER was stained with the ER-ID Red Assay Kit and its total intensity measured following exposure to both drugs, as well as after exposure to DTT as a negative control. Importantly, HU treatment led to a clear increase in ER total intensity, which is an indicative of an increment in ER size^55,56^, while DTT treatment reduced its intensity. This shows that HU-induced ER expansion is conserved in eukaryotic cells.

**Supplementary figure 5.**
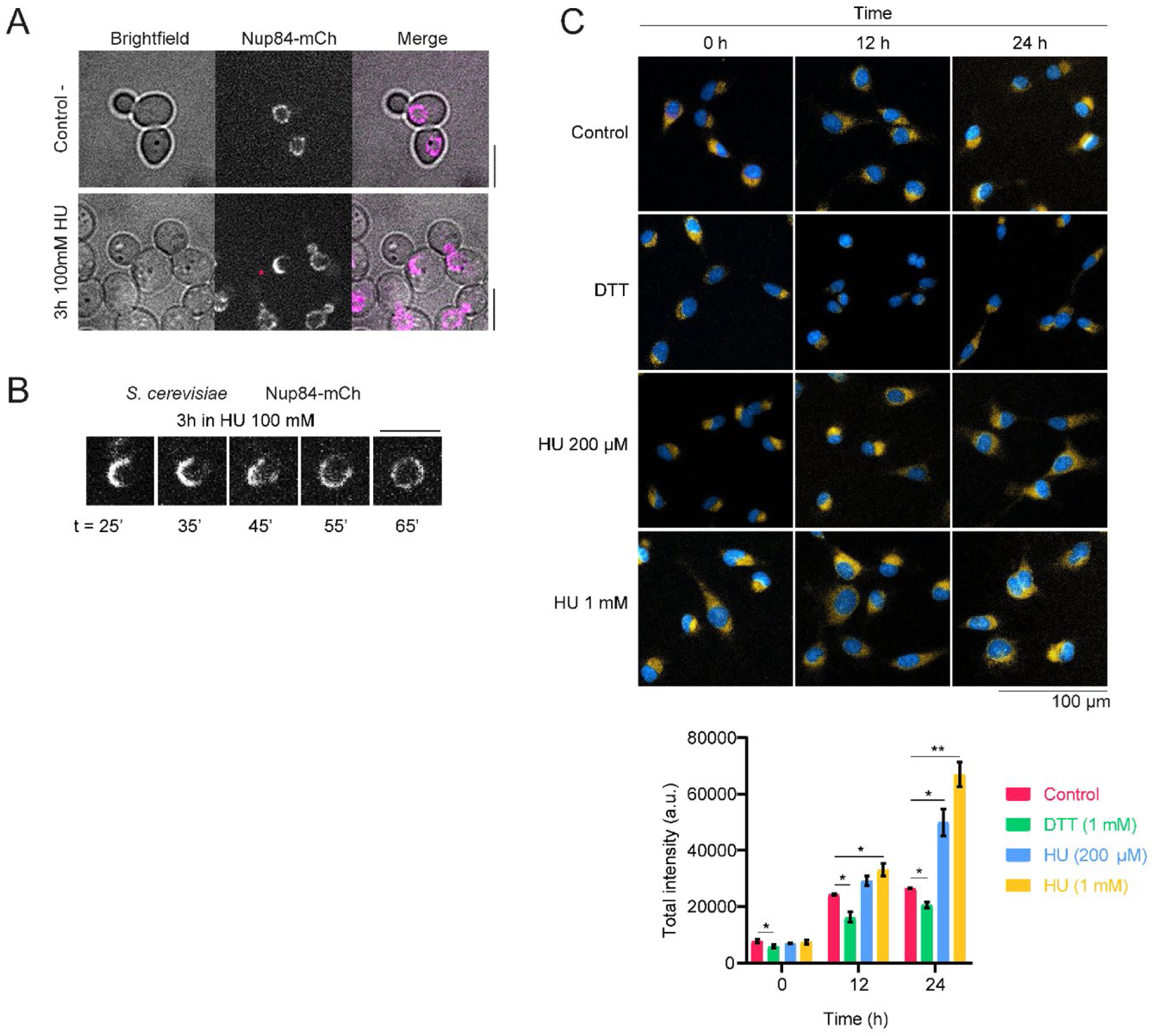
HU-induced perinuclear architecture alterations are evolutionarily conserved. **(A)** Comparison of *S. cerevisiae* cells expressing mCherry-tagged NUP84 in control conditions (left) and after 3 hours in HU 100mM (right). The asterisk marks a cell with clustered NPCs. Confocal microscopy images are SUM projections of three central Z slices. Scale bar represents 5 μm. **(B)** In a *S. cerevisiae* nucleus tagged with NUP84-mCherry, NPCs eventually recover their even distribution along the NE following drug washout after a 3-hour incubation in 100 mM HU. t = time after HU washout. Confocal microscopy images are SUM projections of three central Z slices. Scale bar represents 5 μm. **(C) Upper panel:** Merge of fluorescence images showing Hoescht 333248 (blue, nucleus) and ER-ID® Red (yellow, endoplasmic reticulum) at 0, 12 and 24 hours post-treatment with HU at 200 μM and 1 mM, DTT 1 mM or the equivalent diluent (control). Scale bars represent 100 μm. **Lower panel:** Cells were monitored at 0, 12, and 24 hours post Hydroxyurea (HU) treatment, with 3 wells and 5 fields per well counted per time condition. Dyes were applied 45 minutes before each measurement. ER intensity was segmented to correct for cytoplasmic and extracellular signals, and normalized by ER area in square micrometers (μm²), represented as ‘Total intensity’ (AU×μm²). The control condition (C) corresponds to the highest concentration of HU diluent (water). The number of cells analyzed per experimental condition: 0h: C (959), DTT (1333), HU 200 μM (1577), HU 1 mM (1461).

### Inhibition of protein synthesis with puromycin suppresses ER remodeling

As oxidative alterations caused by HU or DIA might be especially critical during protein synthesis, we tested whether inhibiting *de novo* protein synthesis with cycloheximide (CHX) prevented ER expansion. For that, cells were treated with either HU or DIA, and with or without 100 μg/ml CHX, a concentration that fully blocks protein synthesis in *S. pombe*^57^. Treating fission yeast with HU+CHX resulted in 24.67±17.50% (n>100) cells with expanded ER after a 4-hour incubation, whereas treatment with HU alone resulted in 90.33±5.03% of cells (n>100) with expanded ER in the same conditions (Figure 3A, left). However, when DIA was combined with CHX, ER expansion was exacerbated compared to cells incubated with DIA alone (Figure 3A, right). To elucidate this conundrum, we decided to test the effect of puromycin (Pm), a drug that also inhibits protein synthesis but liberates nascent peptides from ribosomes after being incorporated into the elongating chains^58,59^, in contrast to CHX, which ‘freezes’ ribosomes during elongation, so peptides remain attached to them^60,61^. Interestingly, Pm in combination with either HU or DIA resulted in the total suppression of ER expansion (Figure 3B). As releasing nascent peptides from ribosomes with Pm fully suppresses ER expansion, while blocking their synthesis at ribosomes with CHX exacerbates this phenotype in DIA, these results suggest that disulfide stress interferes with peptide folding during their synthesis and translocation into the ER lumen, and ER expansion may constitute a response mechanism to increase ER folding capacity in these conditions. This is consistent with previous studies, which propose that enlargement of the ER lumen constitutes a survival mechanism against ER stress, as it allows more space to contain damaged proteins and time to refold or eliminate terminally unfolded proteins^15,62–67^.

**Figure 3.**
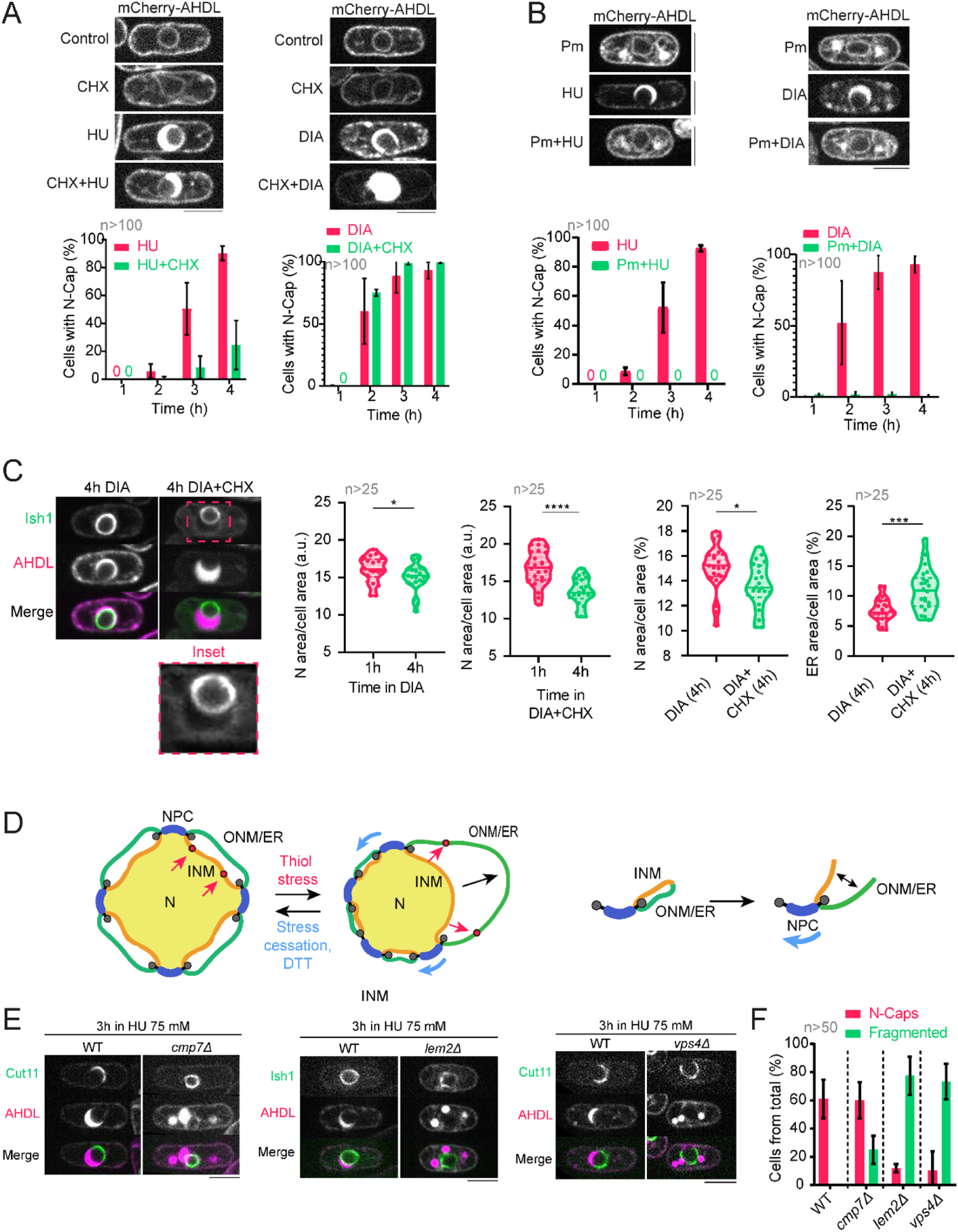
Perinuclear ER expansion is linked to protein synthesis and nuclear size homeostasis. **(A) Upper panels:** Confocal microscopy images of representative cells expressing luminal ER tag mCherry-AHDL exposed to 75 mM HU, 3 mM DIA, 100 μg/mL cycloheximide (CHX) or a combination of the drugs, as indicated, for 4 hours. **Lower panels:** Graphs showing N-Cap incidence in cells exposed to the addressed drugs. **(B) Upper panels:** Confocal microscopy images of representative cells expressing mCherry-AHDL exposed to 75 mM HU, 3 mM DIA, 5 mM puromycin (Pm) or a combination of the drugs, as indicated, for 4 hours. **Lower panels:** Graphs showing N-Cap incidence in cells exposed to the addressed drugs. **(C) Left panels:** Confocal microscopy images of a strain expressing NE tag Ish1-GFP and luminal ER tag mCherry-AHDL exposed to either DIA or DIA combined with CHX for 4 hours. Inset of the N-Cap in DIA+CHX shows a small nucleus opposed to an overextended ER lumen surrounded by Ish1-GFP. **Right panels:** Quantification of the area of either the nucleus (N) or the ER over the area of the whole cell, comparing cells exposed to DIA or to DIA+CHX for 1 and 4 hours. Graphs show violin plots with the mean ± SD of the indicated area coefficient of at least 20 cells in each condition. **(D)** Proposed model for membrane pulling and NPC redistribution during ER expansion. This model hypothesizes a decrease in nuclear size due to the excessive expansion of the ER lumen, which in turn pulls from the INM and forces NPCs to accommodate in the area where the expanded lumen is absent. **(E)** Confocal microscopy images of cells expressing mCherry-AHDL and either Cut11-GFP or Ish1-GFP, comparing a wild-type strain with N-Caps to mutant strains *cmp7Δ*, *lem2Δ* or *vps4Δ*, all of which show a phenotype of fragmented ER, after treatment with 75 mM HU for 3 hours. **(F)** Quantification of the incidence of N-Caps and fragmented ER phenotypes in the total population of cells in the previously addressed conditions. Graphs show the mean ± SD of at least 50 cells in two independent repetitions of the experiment. All confocal microscopy images are SUM projections of three central Z slices. Scale bars represent 5 microns. Graphs in (A) and (B) show the mean ± SD of at least 100 representative cells in each condition in three independent repetitions of the experiment.

### Perinuclear ER expansion is linked to nuclear size homeostasis and requires an intact NE

We noticed that in the presence of DIA and CHX, where perinuclear ER expansion is exacerbated, the size of nuclei was decreased compared to control cells (Figure 3C). This data shows that perinuclear ER expansion has an impact on nuclear size. We hypothesize that, as the perinuclear ER-ONM membrane is continuous with the INM at NPCs, perinuclear ER expansion might retrieve membrane from the INM, especially in the presence of DIA+CHX, a condition in which available membranes might become limiting, resulting in a reduction in nuclear size (Figure 3D).

NE sealing and repair mechanisms rely on the ESCRT-III complex, composed by Cmp7, Vps4 and nuclear specific adaptor Lem2, among other elements. This complex assists with the elimination of aged and damaged NPCs, and repairs and seals discontinuities produced on the NE^68,69^. In addition, Lem2 functions as a barrier to maintain nuclear membrane identity regulating membrane flow between NE and ER^70,71^. Thus, we speculated that, if perinuclear ER expansion is a response to contain cellular damage in the perinuclear ER, then perturbing the NE-ER boundary and NE sealing and repair mechanisms, might result in organelle fragmentation. As expected, addition of HU to cells deleted for *lem2*, *cmp7* or *vps4* resulted in perinuclear ER fragmentation (Figure 3E-F). Together, these data shows that perinuclear ER lumen expansion requires ER-NE remodeling, as well as NE sealing and repair mechanisms to ensure that this expansion occurs while maintaining nuclear and ER compartmentalization.

### Chaperone Bip1 accumulates at the enlarged ER lumen

Bip1 is a Hsp70 chaperone that localizes in the ER luminal space to assists with the folding and translocation of proteins^72–78^. Additionally, a disaggregase activity of mammalian BiP has been recently reported on ER protein aggregates^79^. Thus, we analyzed the localization of Bip1-GFP (kindly provided by M. H. Valdivieso, IBFG) in untreated cells and in cells treated with HU or DIA. In untreated cells, Bip1-GFP localized at the thin perinuclear ER luminal space, as well as in a dotted pattern along the cortical ER (Figure 4A). However, upon HU or DIA treatments, Bip1-GFP cortical foci decreased their number and intensity concomitantly with an increase of Bip1-GFP signal at the expanded perinuclear ER (Figure 4A), whereas the protein levels of Bip1 did not significantly change (Figure 4B). This suggests that most of the cellular pool of Bip1 rapidly relocates from the cortical ER to the expanded perinuclear ER lumen upon ER stress (both in HU or DIA treatments). Our results are in agreement with previous findings in animal cells, where Bip1 is mobile throughout the ER and becomes immobilized upon binding to misfolded substrates^80^. Thus, HU and DIA treatments likely trigger protein unfolding in the perinuclear ER lumen, which mobilizes and retains free Bip1.

**Figure 4.**
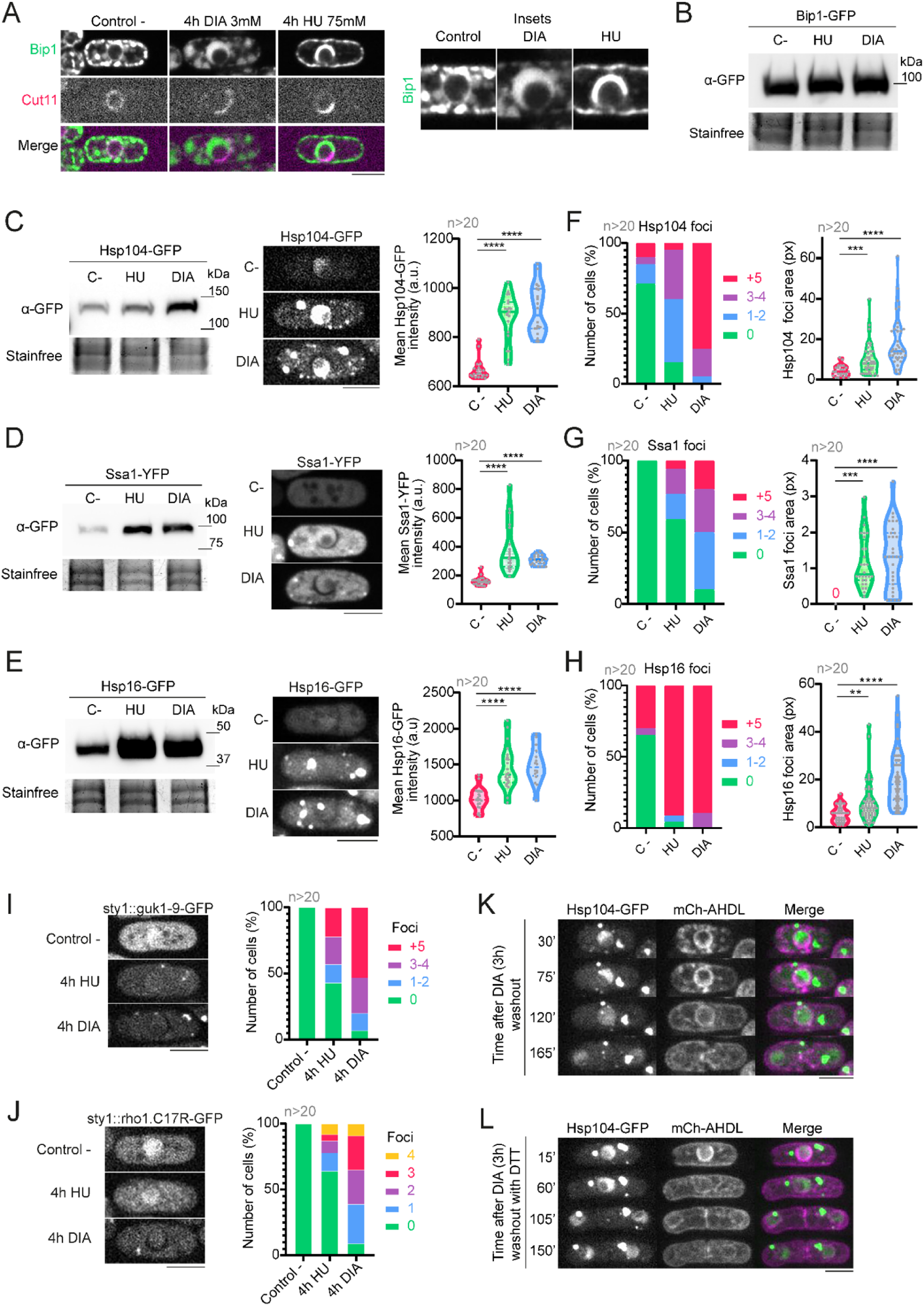
HU and DIA induce cytoplasmic protein and chaperone aggregation. **(A)** Confocal microscopy images showing the distribution of luminal ER chaperone Bip1-GFP in untreated conditions, where it decorates both the perinuclear and cortical ER, and after treatments with 75 mM HU or 3 mM DIA for 4 hours. Bip1-GFP localizes to the expanded perinuclear ER lumen in both conditions, depriving the cortical ER of some of its fluorescence. Panels on the right show insets of the perinuclear region of each cell. **(B)** Western blot showing the total amount of Bip1-GFP, as tagged by an anti-GFP antibody, in control untreated conditions, and after a 3-hour incubation either in 75 mM HU or 3 mM DIA. Bip1-GFP amount remains constant in both treatments as compared with the control. Stain-free lanes are shown as loading control in the lower panels. **(C-E) Left panels:** Western blot showing the total amount of Hsp104-GFP (C), Ssa1-YFP (D) or Hsp16-GFP (E), as tagged by an anti-GFP antibody, in control untreated conditions, and after a 3-hour incubation either in 75 mM HU or 3 mM DIA. All three chaperones increment their total amount in both oxidants with respect to untreated cells. Stain-free lanes are shown as loading control in the lower panels. **Central panels:** Confocal microscopy images of representative cells in control conditions and after 75 mM HU and 3 mM DIA treatments in strains expressing chaperones Hsp104-GFP (C), Ssa1-YFP (D) or Hsp16-GFP (E). In the case of Hsp104-GFP and Hsp16-GFP, cells were incubated for 3 hours in each drug, while cells expressing Ssa1-YFP were incubated for 4 hours. **Right graphs:** Quantification of the total mean intensity of each of the forementioned chaperones in the conditions previously addressed. Graphs of fluorescence intensity (measured in arbitrary units) show violin plots with their respective mean and SD, where at least 20 cells (n) were accounted for in each condition. **(F-H) Left panels:** Quantifications of the number of cytoplasmic foci per cell of Hsp104-GFP (F), Ssa1-YFP (G) or Hsp16-GFP (H) in the conditions addressed in (B-D). Graphs are normalized to the total number of cells per condition, and >20 cells were analyzed for each. **Right panels:** Quantifications of the foci area of the cytoplasmic aggregates noticed with the respective chaperones in the conditions previously addressed. Graphs show violin plots with their respective mean and SD of the foci area (measured in pixels), where at least 20 cells (n) where accounted for in each condition. **(I-J) Left:** Representative cells expressing aggregation markers sty1::guk1-9-GFP (I) and sty1::rho1.C17R-GFP (J), showing foci after a 4-hour incubation in either 75 mM HU or 3 mM DIA. **Right:** Quantification of the number of foci of each of the forementioned aggregation markers in control conditions and after incubation with the two drugs. Graphs are normalized to the total number of cells per condition, and >20 cells were analyzed for each. **(K)** Timelapse showing that cytoplasmic aggregates observed in Hsp104-GFP after DIA treatment, in cells also expressing mCherry-AHDL, do not dissolve over time. **(L)** Timelapse showing that cytoplasmic aggregates observed in Hsp104-GFP after DIA treatment, in cells also expressing mCherry-AHDL, are neither dissolved by 1 mM DTT addition. Confocal microscopy images in (A) are SUM projections of three central Z slices, while confocal microscopy images in (C), (D), (E), (I), (J), (K) and (L) are SUM projections of 18 Z slices. Scale bars represent 5 μm.

### HU and DIA elicit a transcriptional response that differs from canonical unfolded protein response

When proteins do not fold correctly in the ER, the UPR is activated. The major UPR initiator is IRE1, an ER transmembrane kinase endoribonuclease that functions as stress sensor in the ER. When activated by the presence of unfolded proteins, Ire1 performs a non-conventional splicing on Hac1/XBP1 mRNA, which results in the production of UPR transcriptional activators^13,14^. However, *S. pombe* lacks a Hac1/XBP1 ortholog and a UPR-dependent transcriptional response. *S. pombe* Ire1 is responsible for the selective decay of mRNAs encoding ER proteins, thus decreasing the ER load during stress. Bip1 mRNA is the only known mRNA that escapes decay; instead, it suffers a truncation in its 3’-UTR that stabilizes it^81^.

We tested whether HU or DIA-induced ER expansion was dependent on the UPR. To this end, we treated the cells with DTT or tunicamycin (Tn), a drug that impinges on protein glycosylation. Both DTT and tunicamycin are ER stressors known to activate the UPR, which results in cortical ER expansion in *S. cerevisiae*^15,62^. In *S. pombe*, DTT or Tn treatments also led to a phenotype of cortical ER expansion; however, neither drug treatment resulted in perinuclear ER expansion (Figure S6A). Moreover, *ire1Δ* cells expanded the perinuclear ER under DIA treatment with the same frequency and timing than wild-type cells (Figure S6B), which shows that Ire1 is not required for this response. Consistently, whereas *ire1Δ* cells were hypersensitive to DTT, they showed a sensitivity to DIA similar to wild-type cells (Figure S6C).

Thus, to characterize transcriptomic changes associated to ER expansion observed under HU or DIA exposure, we performed an RNA-Seq analysis from cells exposed to each drug at two time points (just prior or during ER expansion; see Methods) as well as untreated wild-type cells. The principal component analysis of the five conditions analyzed pointed to well differentiated transcriptomic programs in DIA and HU (Figure S6D). This was further confirmed in the expression profiles of the upregulated and downregulated genes from both conditions relative to untreated cells (Figure S6E), with waves of overexpression and repression that clearly showed the different patterns between conditions. Notwithstanding, there were 97 upregulated genes (FC ≥ 1, p-value ≤ 0.05) common to HU and DIA treatments when both time points were pooled together, some of which are highlighted in Figure S6F, and 24 that are common to all four individual conditions (Figure S6G). The functional enrichments for the biological processes (Figure S6H) and molecular functions (Figure S6I) of the common transcriptomic signature of both drugs show an upregulation of oxidative stress genes and genes related to iron ion homeostasis. Importantly, chaperones such as Hsp70-Ssa1 and Hsp16, and disaggregase Hsp104 were upregulated relative to control cells, which is an indicative of folding stress. All this considered, the transcriptome analysis under both drugs supports that this ER stress response differs from a canonical UPR response, as many of the genes known to be induced by the UPR, such as Ero11/12, Pdi1 and Ish1 (from their *S. cerevisiae* orthologs stipulated by Travers *et al.*^82^), did not show noticeable alterations in DIA or HU transcriptomes.

Altogether, these results show that ER expansion induced by HU or DIA triggers a transcriptomic response that shares some similarities with the canonical ER stress response, including an upregulation of heat shock response genes, which is an indicative of protein folding stress, but presents a distinct transcriptomic signature of oxidative stress and iron ion unbalance.

**Supplementary figure 6.**
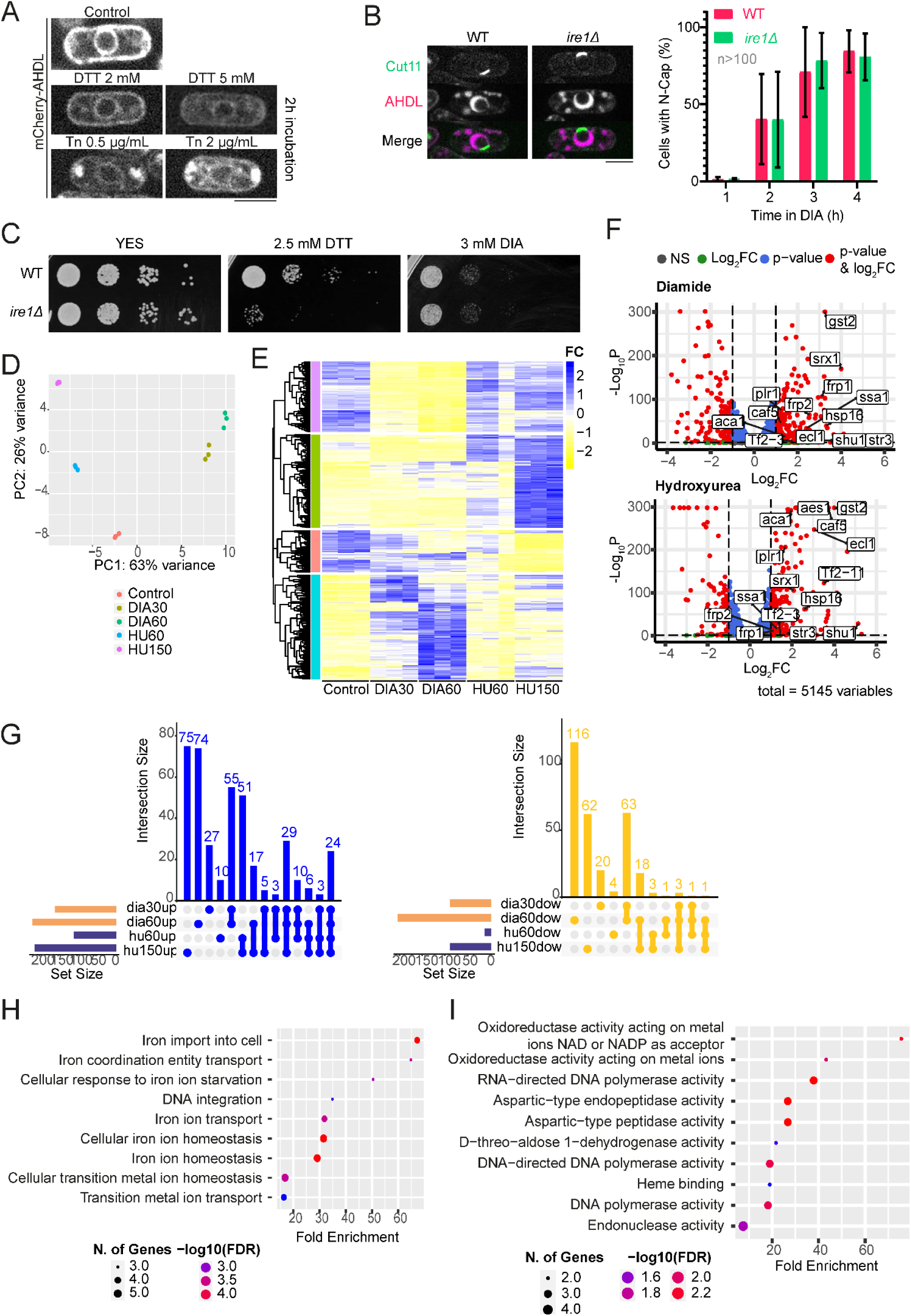
HU and DIA induce a UPR-independent response. **(A)** Canonical ER stress as seen with the luminal ER tag mCherry-AHDL in a wild-type strain after treatment with DTT 2 mM and 5 mM, and with tunicamycin (Tn) 0.6 μg/mL and 2 μg/mL, after a 2-hour incubation. Images are SUM projections of three central Z slices. Scale bars represent 5 microns. **(B) Left panels:** Confocal microscopy images of a representative cell of a wild-type strain and an *ire1Δ* strain, both expressing Cut11-GFP and mCherry-AHDL and showing N-Caps after incubation in 3 mM DIA for 4 hours. Images are SUM projections of three central Z slices. Scale bars represent 5 microns. **Right panels:** Quantification of the incidence of the N-Cap phenotype in a wild-type strain and an *ire1Δ* strain during 3 mM DIA treatment. Graph shows the mean ± SD of three independent repetitions of the experiment, and in each repetition at least 100 cells were accounted for each condition. **(C)** Viability assay of a wild-type strain and an *ire1Δ* strain in control conditions, in presence of 2.5 mM DTT and in 3 mM DIA after a 2-day incubation at 30°C. **(D)** Principal component analysis separating the three independent repetitions of each of the five conditions addressed in the RNA-Seq. Replicates are very consistent, and HU and DIA conditions are well-differentiated between them and with respect to the control. **(E)** Heatmap with the expression profile of upregulated and downregulated genes in 3 mM DIA after 30 and 60 minutes, and in 75 mM HU after 60 and 150 minutes, with respect to an untreated culture. Three replicates of each condition were assessed. Blue indicates upregulation; yellow indicates downregulation; FC = fold change. **(F)** Volcano plots detailing the significantly upregulated and downregulated genes (FC≥1, p-value≤0.05) in DIA (upper graph) and HU (lower graph), pooling together both times addressed. Red dots show genes that surpass the set thresholds of FC and p-value. Highlighted genes are some of the common genes in all conditions; ‘variables’ stands for the total number of genes. **(G)** Upset plots for comparing genes that are upregulated (upper graph) and downregulated (lower graph) between the four treatment conditions. The numbers indicate the genes that are common to the designated combination of conditions. In the case of the unconnected dots, numbers show genes exclusive to that condition alone. Set size graphs indicate the total number of genes in each condition. **(H)** GO term enrichment for the biological processes shared among the genes common to the four treatment conditions. **(I)** GO term enrichment for the molecular functions shared among the genes common to the four treatment conditions.

### HU and DIA induce the accumulation of heat shock response chaperones in cytoplasmic foci

We next analyzed the levels and localization of the chaperones that showed a transcriptomic upregulation upon DIA and HU treatments by fluorescence microscopy. We found that both treatments led to the upregulation of Hsp104-GFP, Hsp70-Ssa1-YFP and Hsp16-GFP relative to control cells, as well as to the presence of cytoplasmic foci, which likely represent protein aggregates (Figure 4C-E). These aggregates increase their number and size over time and are always more prominent in DIA (Figure 4F-H). Consistently, Guk1-9-GFP and Rho1.C17R-GFP reporters of protein misfolding^83^ also accumulated in similar cytoplasmic foci upon DIA and HU treatments (Figure 4I-J). Therefore, these results show that HU and DIA treatments cause defects in protein folding, which not only lead to the expansion of the perinuclear ER and Bip1 relocalization to the expanded luminal space, but also to the accumulation of misfolded proteins and chaperones in the cytoplasm. Interestingly, these cytoplasmic aggregates are not reversible (Figure 4K) nor eliminated upon DTT treatment (Figure 4L), which suggests that accumulated misfolded proteins in the cytoplasm arise independently of ER expansion. As iron ion metabolism seems to be altered upon HU or DIA treatment, as suggested by our RNA-Seq analysis, it is possible that proteins containing conjugated iron ions fail to fold properly under thiol-related stress, giving rise to these terminally unfolded proteins found in cytoplasmic aggregates.

### ER expansion Is GSH-Dependent

DIA forms an intermediate with GSH and induces protein glutathionylation^49,84^. Protein glutathionylation can also be induced by high levels of GSSG, which is in excess in the presence of DIA^85–87^. Alternatively, the absence of GSH upon its sequestration by DIA might impair the vigilance mechanisms that correct, along with PDIs, non-native disulfide bonds in proteins during their folding in the ER^88–90^. Both deregulated glutathionylation and non-native disulfide bond formation lead to protein misfolding^91–94^, which is consistent with the observed accumulation of Bip1 in the enlarged perinuclear ER lumen in both HU and DIA treatments. To test whether GSH plays a role in ER expansion, we manipulated intracellular GSH levels both genetically and chemically. First, we used strains with a deletion of the *gcs1* gene, which encodes glutamyl cysteine ligase, enzyme that catalyzes the rate-limiting initial step in GSH synthesis (union of L-cysteine and L-glutamate into L-γ-glutamylcysteine), and a deletion of the *gsa1* gene, which encodes glutathione synthase, enzyme that catalyzes the final step of GSH biosynthesis (addition of glycine to L-γ-glutamylcysteine)^95,96^, and exposed them to DIA or HU (Figure 5A-B). Importantly, we found that perinuclear ER expansion was suppressed in the absence of Gcs1, and the *gsa1Δ* mutant strain also fully suppressed N-Cap appearance in DIA and significatively reduced its frequency in HU. Moreover, diethyl maleate (DEM), a compound that chemically depletes the GSH cellular pool^97^, also prevented ER expansion in HU or DIA (Figure 5C). Furthermore, when cells were exposed to either HU or DIA in the presence of extracellular GSSG, the phenotype of ER expansion was temporally advanced and significantly increased its incidence (Figure 5D). However, addition of GSSG alone does not induce ER expansion (Figure 5E), indicating that the decrease in the GSH/GSSG ratio is not the trigger of the phenotype, although an imbalance in this equilibrium is crucial for DIA or HU to induce ER expansion.

**Figure 5.**
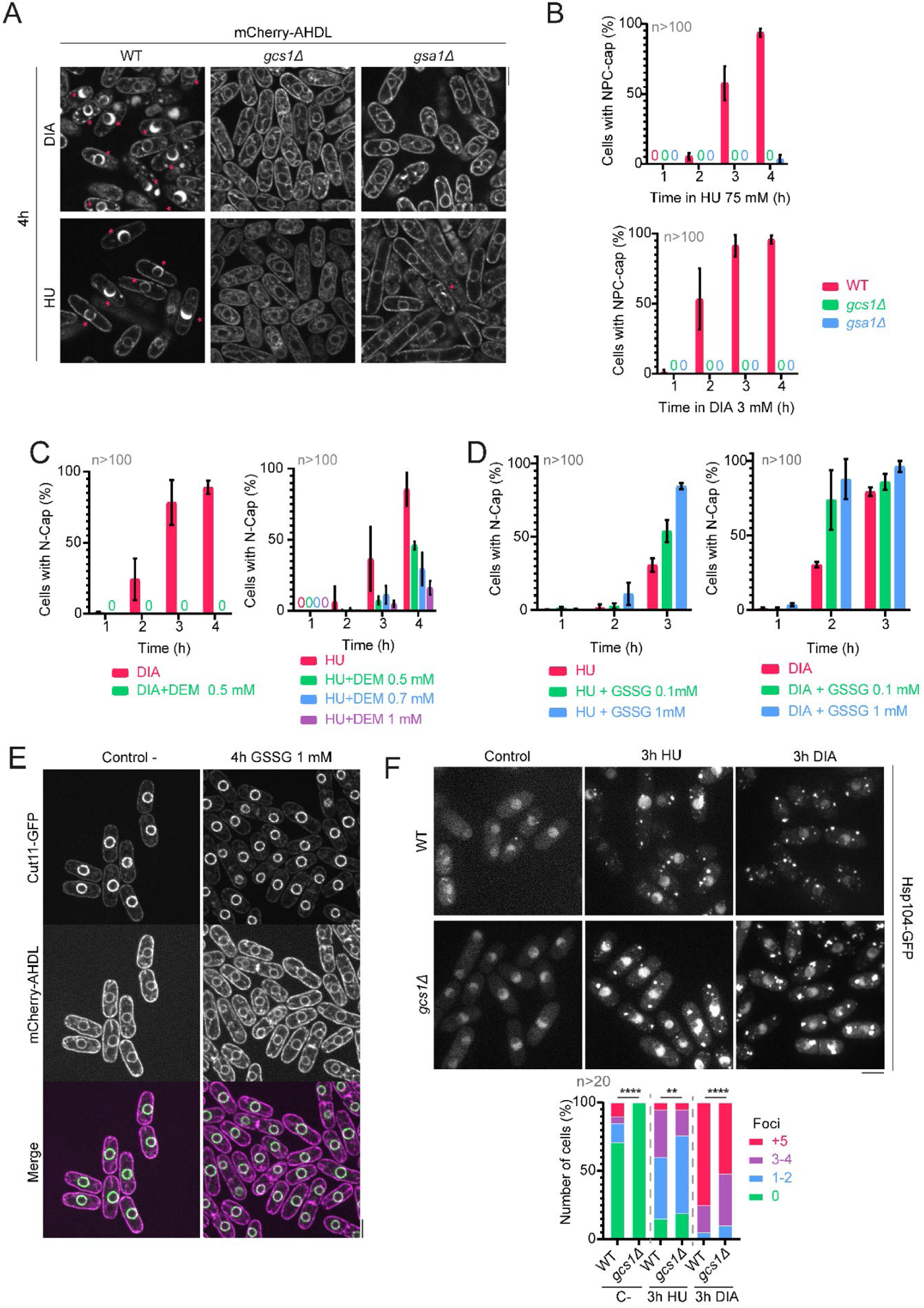
ER expansion is GSH-dependent. **(A)** Confocal microscopy images of wild-type, *gcs1Δ* and *gsa1Δ* strains expressing mCherry-AHDL after a 4-hour incubation in either 3 mM DIA or 75 mM HU. Magenta asterisks indicate cells with N-Caps. **(B)** Quantification of the incidence of the N-Cap phenotype in the previously addressed strains during 75 mM HU (left graph) or 3 mM DIA (right graph) treatments. **(C)** Quantification of the incidence of the N-Cap phenotype in a wild-type strain treated with 3 mM DIA or DIA combined with 0.5 mM DEM (left graph), and with 75 mM HU or HU combined with 0.5 mM, 0.7 mM or 1 mM of DEM (right graph). **(D)** Quantification of the incidence of the N-Cap phenotype in a wild-type strain treated with 75 mM HU or HU combined with 0.1 mM or 1 mM of GSSG (left graph), and with 3 mM DIA or DIA combined with 0.1 mM or 1 mM of GSSG (left graph). **(E)** Confocal microscopy images of wild-type cells expressing Cut11-GFP and mCherry-AHDL in untreated conditions and after a 4-hour incubation in the presence of 1 mM GSSG. The internal morphology of the cells remains unaltered after exposure to GSSG. **(F) Upper panels:** Confocal microscopy images of wild-type and *gcs1Δ* strains expressing Hsp104-GFP after a 3-hour incubation in either 3 mM DIA or 75 mM HU. Images are SUM projections of three central Z slices. Scale bars represent 5 microns. **Lower panels:** Quantification of the number of Hsp104-GFP foci observed in a wild-type and a *gcs1Δ* strain after a 3-hour treatment with either HU or DIA. The graph is normalized to the total number of cells per condition, and >20 cells (n) were analyzed for each. Confocal microscopy images in (A) and (E) are SUM projections of three central Z slices, while confocal microscopy images in (F) are SUM projections of 18 Z slices. Scale bars represent 5 μm. Graphs in (B), (C) and (D) show the mean ± SD of two independent repetitions of the experiment, and in each repetition at least 100 cells were accounted for each condition.

Finally, we determined that deletion of *gcs1* did not suppress the accumulation of Hsp104-GFP in cytoplasmic foci (Figure 5F), which further supports that cytoplasmic protein aggregates induced by HU and DIA are independent from ER expansion and arise from an imbalance in the cytoplasm that is not a direct consequence of thiol stress, as it is not reverted by DTT addition (Figure 4L). Whether this is due to the different GSH/GSSG ratios in the ER and cytoplasm, or if these aggregates emerge as a consequence of an iron ion imbalance would require further investigation.

## DISCUSSION

In this work, we show that HU causes ER stress characterized by a remarkable expansion of the perinuclear ER lumen that accumulates Bip1, as well as cytoplasmic aggregation of proteins and chaperones, suggesting that HU interferes with protein folding in both compartments. HU-induced ER expansion is independent of the inhibition of RNR activity, the best-known target of HU, as well as of S phase arrest, and is recapitulated by thiol stress inductor DIA. Puromycin, which blocks protein synthesis and releases nascent peptides from ribosomes, suppresses HU- or DIA-induced ER expansion. DTT, a strong reductant, both prevents and suppresses ER remodeling caused by both drugs, although it cannot revert cytoplasmic aggregate formation. Thus, based on these observations, we hypothesize that HU causes imbalances in the cell redox homeostasis, which in turn results in cytoplasmic and ER protein misfolding. Importantly, ER stress induced by HU involves GSH, as interfering with its synthesis or sequestering it chemically prevents ER expansion, the same which occurs with DIA. Thus, like DIA, HU causes protein unfolding in a GSH-dependent manner.

Our results show that ER expansion involves INM and ONM-ER dissociation and leads to NPCs clustering along with the SPB in one region of the NE opposite to the expanded lumen. NPC clustering has been mostly associated with aged or defective NPCs and also observed in pathological conditions^98^. Despite demonstrating that clustered NPCs remain functional for nucleocytoplasmic transport and that ER expansion is a transient event, biological consequences of a clustered NPC-SPB architecture in genome organization, gene transcription, and NE epigenetic environment remain an open question.

Nuclear and ER size homeostasis is tightly linked to the regulation of membrane flow between the ER and the nucleus^4,70,99^. This flow is highlighted in our study by two independent observations: first, defective NE sealing and repair results in ER fragmentation upon its expansion, and second, the reduction of nuclear size in conditions of exacerbated ER expansion. This reduction of nuclear size might directly impact nuclear protein concentration, whose biological consequences will require further investigation.

Our observations in the fission yeast regarding ER expansion are consistent with previous works reporting that perturbations in its homeostasis can result in expansion of ER membranes and luminal volume to increase its protein folding capacity, preventing unfolded or misfolded protein from forming aggregates^15,62,100–103^. Moreover, ER swelling has been demonstrated to be reversible under ER stress conditions^103–109^, and cortical ER expansions induced by DTT or Tn have been shown in budding yeast, a response which requires activation of the UPR^15^. However, HU- or DIA-induced ER expansion occurs in the absence of *ire1,* and transcriptomic analysis of HU- or DIA-treated cells does not reflect an upregulation of lipid synthesis nor other genes associated with the UPR. This supports the hypothesis that ER remodeling observed upon HU or DIA treatment is UPR-independent.

According to our results, it has been proposed, in works done largely in *C. elegans*^110^ and in *S. cerevisiae*^111^, that oxidative stress is able to inhibit the canonical signaling pathway of Ire1, which sets a precedent in UPR-independent ER stress responses. In fact, *S. pombe* lacks a canonical UPR-dependent transcriptional response^81^. Therefore, as previously described in conditions of ER stress, HU- and DIA-induced ER expansion might constitute a UPR-independent proteostatic mechanism to favor protein folding, specifically under thiol-related stress, by providing extra luminal space in an organelle highly sensitive to redox perturbances.

We have shown that HU and DIA exposures also lead to the formation of cytoplasmic aggregates. Our transcriptomic data under HU or DIA revealed a deregulation of iron ion homeostasis. Iron balance is essential for the proper folding of some proteins and enzymes^112,113^, and HU has been described to have a deleterious effect on cytosolic Fe-S clusters in yeasts^114^. Interestingly, this effect seems to be indirect, as it is observed *in vivo* but not *in vitro*^114^, and GSH is required for the maturation of Fe-S clusters^115^. Thus, another possibility could be that the effect of DIA and potentially of HU on the GSH cellular pool might result in defective synthesis and/or folding of iron-containing proteins, leading to their irreversible cytoplasmic aggregation.

In summary, our data strongly that HU deregulates the cell redox homeostasis and results in protein misfolding. This leads to protein and chaperone aggregation in the cytoplasm and in a GSH-dependent perinuclear ER expansion, phenotypes that are recapitulated by DIA. The mechanisms by which HU and DIA achieve the same phenotype remain unclear and likely differ, as DIA directly affects GSH/GSSG pools, while HU might indirectly achieve the same by unbalancing the overall redox equilibrium or by tampering with ion homeostasis. Notwithstanding, our results show unprecedented cellular alterations induced by HU tightly related to imbalances in oxidative protein folding.

## ACKNOWLEDGEMENTS

We would like to thank Daga’s lab members and our colleagues the CABD and Genetic Area for useful discussions. We thank Victor M. Carranco (Genetics area), Katherina García (CABD microscopy facility) and Laura Tomás (CABD proteomic facility), for their excellent technical help. High-Pressure Freezing, Freeze Substitution and TEM experiments has been performed by the ICTS “NANBIOSIS”, more specifically by the U28-Nanoimaging Unit at IBIMA Plataforma BIONAND. We also thank Henar Valdivieso, Paul Nurse, Snezhana Oliferenko, Olaf Nielsen, Fred Chang, Peter Walter, Ralf E. Wellinger, and the Yeast National Bioresource Project (Japan) for kindly sharing strains. We thank C3UPO for the HPC support. This work was supported by the *Ministerio de Economía y Competitividad* from the Spanish government (grant: PID2021-128408OB-I00 to RRD). ASM is founded by a FPU fellowship (FPU18/02016). MB was supported by A.4. Fellowship *(“Ayudas para la Incorporación de Doctores”*), II Plan Propio (University of Malaga).

## METHODS

### Culture conditions

Standard methods for *S. pombe* growth and genetics were used^116^. Liquid cultures were grown in YES medium at 30°C in shaking incubators. Pre-inoculations of these cultures were left growing overnight and diluted the next day, allowing the culture to grow up to mid-log phase. Mating was performed on SPA solid medium plates and the spores generated in the crosses were dissected using a dissection microscope (MSM 400; Singer Instruments) on YES agar plates.

### Strain construction

*S. pombe* strains used in this work are listed in Table S1. Combination of GFP-, mCherry- or tdTomato-tagged proteins and deletion mutants were generated by mating strains already available in our laboratory or shared with us by other groups and posterior tetrad analyses. The strains used in this study were derived from original strains provided by F. Chang (University of California, San Francisco, CA), T. Toda (Graduate School of Integrated Sciences for Life, Hiroshima University, Higashi-Hiroshima, Hiroshima, Japan), J. Jiménez and V. Alvarez (Centro Andaluz de Biología del Desarrollo/Universidad Pablo de Olavide, Seville, Spain), P. Nurse (The Francis Crick Institute, London, UK), S. Moreno (Instituto de Biología Funcional y Genómica, Salamanca, Spain), Y. Watanabe (Institute of Molecular and Cellular Biosciences, University of Tokyo, Tokyo, Japan), I. Hagan (Cancer Research UK, University of Manchester, Manchester, UK), Y. Hiraoka (Advanced ICT Research Institute, Kobe, Japan; National Institute of Information and Communications Technology, Kobe, Japan/Graduate School of Frontier Biosciences, Osaka University, Suita, Japan), O. Niwa (Kazusa DNA Research Institute, Chiba, Japan), S. Oliferenko (The Francis Crick Institute, London, UK), M. H. Valdivieso Montero (Instituto de Biología Funcional y Genómica, Salamanca, Spain) and the National Bio-Resource Project (Osaka, Japan).

**Table S1.** Strains used in this work.

| Number | Genotype |
| --- | --- |
| **RD257** | h- cdc25-22 cut11-GFP:ura |
| **RD1936** | h+ wild-type U-L- |
| **RD4595** | h+ pBip1-mCherry-AHDL::leu1 ura4-D18 ade6- |
| **RD4596** | h- pBip1-mCherry-AHDL::leu1 ura4-D18 ade6- |
| **RD6766** | h sid2-tomato-nat cut11-GFP-ura |
| **RD6911** | h+ pBip1-mcherry-AHDL::leu1 cut11-GFP-ura |
| **RD6912** | h- pBip1-mcherry- AHDL::leu1 cut11-GFP-ura |
| **RD6990** | h cut11-GFP-ura nup211-mcherry-nat |
| **RD7174** | h pBip1-mCherry-ADHL::leu1 ish1-GFP-Kan |
| **RD7255** | h pBip1-mcherry-AHDL::leu1 imp1-GFP-Kan |
| **RD7555** | h heh1(lem2)-GFP-Kan pBip1-mcherry-ADHL::leu1 |
| **RD7733** | h+ lem2::ura4+ ish1-GFP-KanMX6 pBip1-mcherry-ADHL::leu1 |
| **RD7735** | h kan-nmt41-GFP-ima1 pBip1-mcherry-ADHL::leu1 |
| **RD7737** | h man1-GFP-Hph pBip1-mcherry-ADHL::leu1 |
| **RD7924** | h nmt41-pap1-GFP-leu nup211-mCherry-Nat |
| **RD7967** | h+ cut11-GFP::ura4+ leu1-32 |
| **RD7968** | h- cut11-GFP::ura4+ leu1-32 |
| **RD7971** | h+ hsp104-GFP-ura pBip1-mcherry-AHDL::leu1 |
| **RD7971** | h+ hsp104-GFP-ura pBip1-mcherry-AHDL::leu1 |
| **RD7999** | h ssa1-YFP-Kan pBip1-mcherry-AHDL::leu1 |
| **RD8018** | h cdc22-M45 cut11-GFP-ura |
| **RD8163** | h spd1::Hph cut11-GFP-ura |
| **RD8428** | [*S. cerevisiae*] nup84-mcherry rad52-GFP |
| **RD8723** | h nup120::nat cut11-GFP-ura pBip1-mcherry-ADHL::leu1 |
| **RD8768** | h cmp7::kan cut11-GFP-ura pBip1-mCherry-AHDL::leu1 |
| **RD8884** | h ire1::kan pBip1-mCherry-AHDL::leu1 cut11-GFP-ura |
| **RD8971** | h vps4::Hph cut11-GFP-ura pBip1-mcherry-AHDL::leu1 |
| **RD9060** | h- hsp16:GFP:NatMX6 |
| **RD9074** | h hsp16:GFP:NatMX6 pBip1-mcherry-AHDL::leu1 |
| **RD9076** | h sty1::guk1-9-GFP::leu1 pBip1-mcherry-AHDL::leu1 |
| **RD9077** | h gcs1::KanMX6 cut11-GFP-ura pBip1-mcherry-AHDL::leu1 |
| **RD9102** | h sty1::rho1.C17R-GFP::leu1 alm1-tomato-nat |
| **RD9135** | h gcs1::kan hsp104-GFP-ura alm1-tomato-nat |
| **RD9243** | h gsa1::kan cut11-GFP-ura pBip1-mCherry-AHDL::leu1 |
| **RD9344** | h bip1-GFP:leu1 cut11-RFP-Hyg |

### Cell synchronization

For synchronization in S phase using hydroxyurea (127-07-1; Sigma-Aldrich), cells were grown in YES media at 30°C up to a OD_600_ of 0.3, and then HU was added from a 1M stock dissolved in YES liquid medium to a final concentration of 15 mM, keeping the culture at 30°C for at least 3 hours. For *cdc25-22* thermosensitive strains (G2 arrest), cells were grown at 25°C up to OD_600_ of 0.3 and then shifted to 36°C for 4 hours.

### Viability assays

Viability assays, or drop assays, were performed by plating 5 μL of a OD_600_=0.3 cell culture of the indicated strain and then plating the same volume of three 1:10 sequential dilutions. Control plates consisted of YES agar and were incubated at 30°C for 2-3 days. The rest of the plates were prepared by melting solid YES agar and adding the indicated drug to the concentration stated in the corresponding figure while the medium was still warm. After cooling, an exact replicate of the control disposition was plated and incubated at 30°C for 2-3 days.

### Drug treatments

Hydroxyurea (127-07-1; Sigma-Aldrich) was used at a final concentration ranging from 15 to 200mM, as indicated in each experiment. Diamide (D3648; Sigma-Aldrich) was added to the sample at a final concentration of 3 mM. DETA-NONOate (A5581; Sigma-Aldrich) was added to the sample at a final concentration of 0.5 mM. Diethyl maleate (D97703; Sigma-Aldrich) was added to the sample at a final concentration of 0.5 mM, 0.7 mM or 1 mM. Dithiothreitol (R0862; Thermo Scientific) was added to the sample at a final concentration of 1, 2 or 5 mM. Tunicamycin (T7765; Sigma-Aldrich) was added to the sample at a final concentration of 0.5 or 2 μg/mL. Cycloheximide (C7698; Sigma-Aldrich) added to the sample at a final concentration of 100 μg/mL. Puromycin (BIP3340; Apollo Scientific) was added to the sample to a final concentration of 5 mM; cultures were pretreated for 30’ with this drug before adding DIA or HU to ensure complete inhibition of protein synthesis. L-Glutathione oxidized (G4501; Sigma-Aldrich) was added to the sample to a final concentration of 1 or 10 mM. Menadione (M5625; Sigma-Aldrich) was added to the sample to a final concentration of 50 μM. Hydrogen peroxide (H1009; Sigma-Aldrich) was added to the sample to a final concentration of 0.2, 1 or 5 mM.

### RNA-Seq and transcriptomics analysis

RNA was extracted from 200 mL cultures grown in agitation at 30°C, then incubated under the desired conditions (untreated cultures [C-]; cultures treated with 75 mM HU for 60 minutes [HU60] and for 150 minutes [HU150]; and cultures treated with 3 mM DIA for 30 minutes [DIA30] and for 60 minutes [DIA60]. HU60 and DIA30 correspond to a time prior N-Cap formation; HU150 and DIA60 mark a time point in which cells begin to show the phenotype). Cells were then pelleted and ruptured using a precooled mortar with liquid nitrogen. After cell lysis, debris was recovered and snap-frozen in liquid nitrogen. RNAase-free water was then added to the pellets and RNA was extracted using the RNAeasy kit (74104; Qiagen). Final RNA concentration was adjusted to 5 ng and sent to BGI for sequencing. The bulk data were then processed using a customized BaSH script. Read assembly was performed using STAR 2.7.10a^117^, mapping was performed on the *S. pombe* genome version ASM294v2.48 using SAMtools 1.3^118^, and the final count matrix was performed with htseq-count from package HTSeq 2.0.1^119^. For the search of differentially expressed genes, a script in the R programming language was used, using Bioconductor 3.19 and the DESeq2 package^120^. The GO term enrichment was performed using DAVID^121^ and ShinyGO^122^.

### *In vivo* measurement of H_2_O_2_ levels with roGFP2-Tpx1.C169S

Auxotrophic strain HM123 (*h^−^ leu1–32*, lab stock) was transformed with plasmid p407.C169S and grown in filtered minimal media at 30°C as previously described^123^. Stationary phase pre-cultures were diluted to reach an OD_600_ of 1 after 4-5 duplications. We determined the degree of sensor oxidation (OxD), as previously described^124^ being 1 the maximum oxidation of the probe reached with 1 mM H_2_O_2_ treatment, and 0 the complete reduction of the biosensor with 50 mM DTT. After recording 4 cycles, treatments were added as following (final concentrations): water as control, 0.1 mM H_2_O_2_, 3 mM of DIA, and 75 mM HU.

### Microscopy and image analysis

Live cell imaging was performed at 25°C, with a Roper Scientific spinning disk confocal microscope (IX-81; Olympus; CoolSnap HQ2 camera, Plan Apochromat 100×, 1.4 NA objective), or a 3i Marianas CSU-W1 spinning disk (50µm pinholes) confocal microscope. Images were acquired and analyzed with ImageJ/Fiji (National Institutes of Health). For imaging, cells were immobilized in soybean lectin (Sigma)-coated μ-Slide 8-well dishes (Ibidi; Cat. No. 80827) in the case of *S. pombe*; immobilization *of S. cerevisiae* was achieved placing an YPD+agar 1% pad upon the cells in the μ-Slide 8-well dishes. Unless otherwise stated, images shown are Z projections of 3 central z sections with a step size of 0.3 μm. Quantifications of total fluorescence intensity were performed on SUM projections of 18 z sections with a step size of 0.3 µm. Roundness measurements were performed in a single central z section. The ‘roundness’ parameter in the ‘Shape Descriptors’ plugin of Fiji/ImageJ was used after applying a threshold to the image in order to select only the more intense regions and subtract background noise^125^. Roundness descriptor follows the function:

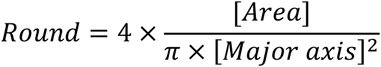

where [*Area*] constitutes the area of an ellipse fitted to the selected region in the image and [*Major axis*] is the diameter of the round shape that in this case would fit the perimeter of the nucleus.

### Transmission Electron microscopy

This protocol, based on previous works^126,127^, was performed using the Leica EM AFS2 device for freeze substitution resin-embedding at low temperatures. *S. pombe* samples were subjected to high pressure freezing using the Leica EM HPM100 system, and then transferred from liquid nitrogen to the AFS2 chamber, where the freeze substitution was initiated. Samples were washed thrice with cold acetone and post-fixed with 1% osmium tetroxide (dissolved in acetone) for 2 hours at 4°C in darkness, after which they were again washed thrice with acetone. Spurr resin (prepared as described by Spurr^128^) inclusion was then initiated as follows: 1:1 resin/acetone overnight at room temperature, 3:1 resin/acetone 4-8 hours at room temperature, 100% resin overnight at room temperature. After that, samples are plated with 100% resin and polymerized for 2-3 days (at least 24 hours) at 70°C. Electron microscopy images were then acquired.

### HT1080 fibrosarcoma cell preparation and cultivation

HT1080 fibrosarcoma cells were cultured in DMEM medium supplemented with 10% fetal bovine serum (FBS; Capricorn Scientific, Ebsdorfergrund, Germany), a combination of 10,000 IU/mL penicillin and 10,000 μg/mL streptomycin (Corning, MA, USA), and 2 mM L-glutamine (Merck, Darmstadt, Germany). The cells were maintained at 37°C in a 5% CO_2_ humidified atmosphere. Cultures were grown in flasks until they reached 70-80% confluence. Subsequently, 8500 cells were seeded per well in a 96-well plate with a final volume of 200 μL and incubated overnight prior to conducting various endoplasmic reticulum assays.

### High Content Screening Analysis of Endoplasmic Reticulum Stress

Images were captured using the Operetta High Content Imaging System (PerkinElmer, Inc.). Images were collected at different time points (0, 12, and 24 hours) from various wells to assess the duration of cell exposure to the compounds. Fluorescent dyes (Hoechst 33342 [H1399; Invitrogen^TM^, Thermo Fisher Scientific], for DNA, added at a working concentration of 2.5 μM/mL; CellMask^TM^ Deep Red [C10046; Invitrogen^TM^, Thermo Fisher Scientific], for the plasma membrane, added at a working concentration of 0.25 μM/mL; ER-ID® Red assay kit [ENZ-51025-K500; Enzo Life Sciences], for the endoplasmic reticulum, added at a working concentration of 1:2000) were uniformly added and incubated for 1 hour before imaging. Image analysis was conducted using Harmony software version 4.8 (PerkinElmer, Inc.). Cells that moved out of the selected field of view or underwent division, either during initial imaging or at later time points, were excluded from the analysis. The signal was corrected by subtracting the cytoplasmic signal (*S_cyto_*) from the ER signal (*S_ER_*), and the external signal around each cell (*S_surround_*) was then subtracted from this adjusted ER signal. This process ensures that the final intensity measurement (*S_corrected_*) accurately reflects the corrected ER signal isolated from both the cytoplasmic and surrounding signals:

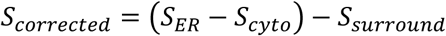

Each well was divided into 4/5 distinct fields of view to ensure comprehensive data collection. The acquired images from the Operetta High Content Imaging System were analyzed and processed using FIJI/ImageJ.

### Quantification and statistical analysis of microscopy images

At least three independent biological replicates were performed for each experiment. Graphs and statistical analyzes were performed with GraphPad Prism 5.0 (GraphPad Software) and Excel (Microsoft). Unless otherwise stated, graphs represent the mean and error bars represent the standard deviation (SD). n is the total number of cells scored in each repetition of the experiment, as described in each figure legend. Statistical comparison between groups in graphs depicting N-Cap incidence and violin plots was performed by unpaired Student’s t test, considering two-tailed p-values exceeding 0.05 to be not significant. Statistical comparison between groups in graphs accounting for number of foci was performed by chi-squared test. Asterisks in graphs correspond to the following p-values: (ns) P > 0.05; (*) P ≤ 0.05; (**) P ≤ 0.01; (***) P ≤ 0.001; (****) P ≤ 0.0001.

